# Nanodisc-Enabled Dissection of GxxxG- Mediated Amyloid Beta Oligomerization Mechanisms

**DOI:** 10.64898/2026.11.13.766326

**Authors:** Dibakar Sarkar, Dipanwita Roy, Karishma Biswas, Satadru Chakraborty, Suprity Majumder, Janarthanan Krishnamoorthy, Nirakar Sahoo, Narayan Maity, Chandan Singh, Dhiman Ghosh, Ranjit Biswas, Dulal Senapati, Prasenjit Bhaumik, Anirban Bhunia

## Abstract

Extracellular deposition of amyloid plaques is a major hallmark in Alzheimer’s disease (AD), in which amyloid-beta (Aβ) fibrils are key constituents. Fibrillation of Aβ is modulated by several key sequence motifs, among which the G^33^LMVG^37^ motif located in the C-terminal is particularly significant. Moreover, the interaction between Aβ and cellular membrane can be determining factor in its oligomerization. Here, we present membrane scaffold protein (MSP) encased lipid nanodiscs as a model membrane mimic to investigate the roles of G33 and G37 in membrane-mediated Aβ40 oligomerization. Using a combination of spectroscopic and microscopic techniques, accompanied by molecular dynamic (MD) simulations, we demonstrate that these Gly residues are crucial in regulating both self-assembly and membrane-associated oligomerization of Aβ40. We report the formation of structurally distinct Aβ40 oligomers upon interaction with nanodiscs. Our mutational studies reveal a direct involvement of G33/37 residues in membrane interaction, leading to prolonged oligomer lifetime.

## 1. INTRODUCTION

Aggregation of amyloid beta (Aβ) peptides in the brain has been implicated as a major hallmark of Alzheimer’s disease (AD)(1). Aβ peptides are produced through the sequential cleavage of the trans membrane amyloid precursor protein (2), (3) which then undergoes self-association along a nucleated growth pathway, with oligomeric intermediates being the primary culprits behind AD-pathogenesis(4),(5). Despite decades of research, the collective discoveries have not yet converged on a single universal mechanism of amyloidogenesis (6),(7). One of the major challenges in understanding the comprehensive mechanisms of amyloidogenesis is the inherent conformational plasticity of these intrinsically disordered proteins(8), which leads to high degree of polymorphism(9) in both fibrillar and pre-fibrillar structures (10),(11). Additionally, cell membrane plays a crucial role in modulating amyloid aggregation (12),(13)and directing aggregates to toxic intermediate states (14,15). Several studies have suggested that Aβ monomers either forms ion-selective pores on cell membranes or cause lipid fragmentation during fibrillation(16). The Aβ-membrane interaction also triggers conformational change (17),(18) in the peptide’s secondary structure, influencing the fibrillation process(19) (20). Several factors like membrane composition, charge, fluidity, and curvature govern the Aβ-membrane interaction (20), (21). In fact, the lipid composition of the bilayer can attenuate aggregation kinetics, leading to structural polymorphisms(22). In a recent study, our group has demonstrated a detailed insight on the impact of different membrane compositions (a simplified blood-brain barrier mimic, composed of POPC/POPG/GM1/ cholesterol, and the native total brain lipid extract) on Aβ40 aggregation dynamics (12) (23). The crucial role played by the hydrophobic interactions between the lipid acyl chains and residues spanning K16-K28 and I31-V36 in forming transient conformations during peptide aggregation was further elucidated. We observed a temporal transition in the secondary structure of Aβ40 towards helical intermediates, highlighting membrane-induced transient folding(12).

Recent studies show that lipid nanodiscs (NDs)(24),(25) are increasingly being used as models of natural cell membranes to study how these membranes affect amyloid aggregation (26), (27,28). Although these phospholipid bilayer NDs don’t exactly match real cell membranes, they have important advantages over older models like micelles or vesicles (27), (29) (30). Typically, NDs are very homogeneous and firmly soluble in a wide range of buffer systems(31). They allow researchers to create specific lipid mixtures and precisely control the size and charge of the bilayer (32), (33). On the other hand, small unilamellar vesicles (SUVs) are known to undergo rapid changes in the lipid environment (from homogeneous to heterogeneous) upon interaction with amyloid proteins (34). Moreover, the increased stability of NDs mediate interaction with a stable planar bilayer while reducing the effect of membrane curvature. Notably, NDs have been previously utilized for studying lipid specificity, as well as for the entrapment of Aβ oligomers (35), (36). Overall, the precise lipid composition embedded within NDs of desired size, coupled with their high stability and homogeneity allows for comprehensive quantitative understanding of Aβ-membrane association.

Over the years, studies have revealed the crucial role played by the central (K16-G25) and the C-terminal (K28-G37) region in Aβ self-assembly and membrane interaction (37), (38) Interestingly, the conserved GxxxG motif present at the C-terminus of Aβ possesses unique functional characteristics throughout the Aβ aggregation pathway, making it a particularly noteworthy target for understanding the physiological and pathological roles of transmembrane glycine zippers in membrane-associated proteins.(39) While this motif has been linked to APP processing and subsequent Aβ production (40), its significant role in regulating Aβ oligomer formation and neurotoxicity is also evident from previous studies (41), (42), (2). In this context, the precise role of the GxxxG repeating motifs in the C-terminal of Aβ peptide fragment in maintaining structural dynamicity and neurotoxicity has been recently demonstrated(43). The GxxxG motif forms an integral part of the transmembrane domain of the peptide, and the Gly residues are particularly involved in imparting flexibility and dynamics to the peptide, thereby influencing membrane insertion through ion-channel formation. This, in turn, contributes to the toxicity of Aβ peptides and leads to neuronal cell death. Kim et al. reported that G → L mutation in the GxxxG motif, particularly G37L in Aβ, decreased ion-channel induction in synthetic membranes and diminished toxicity in Neuro 2a neuroblastoma cells (41). Hung et al. also reported that G → L mutants of Aβ42, the GSL peptides (G25L, G29L, G33L, or G37L), showed an increased rate of amyloid formation and a decrease in the formation of toxic oligomeric species in a murine model (2). Our group has also demonstrated that the GxxxG motifs in the C-terminal Aβ fragment (A21–V40) are actively involved in preserving structural flexibility and neurotoxicity. Substitution of key Gly residues alters the hydrogen-bonding network, thereby affecting helix–helix interactions and leading to distinct fibrillation pathways. Specifically, replacing Gly with Leu within the C-terminal GxxxG motif markedly accelerates aggregation kinetics. Notably, when the G→L substitution is positioned closer to the C-terminus, aggregate-size heterogeneity decreases significantly. These results underscore the critical role of Gly residues in regulating both aggregation rates and the distribution of aggregate sizes (43).

In addition, cytotoxicity toward neuronal SHSY5Y cells declines substantially as the Leu substitution approaches the C-terminus. The GxxxG motif appears to promote greater conformational heterogeneity, which correlates with increased cytotoxicity, whereas its mutation facilitates a rapid transition from random coil to β-sheet structure. Hydrogen-bonding networks are central to conformational selection, and Leu substitution disrupts these interactions, destabilizing intermediate conformers and thereby reducing neurotoxicity. These findings align with the growing emphasis on prefibrillar species as key contributors to disease pathophysiology (43), (44).

In this study, we investigate the significance of GxxxG motif in the membrane-mediated oligomerization of Aβ40, in a near-native membrane environment. To elucidate the role of the GxxxG motif, and in particular the contributions of residues G33 and G37 to membrane-mediated oligomerization of Aβ40, a set of peptides was designed (Figure 1A), including the full-length Aβ40, Aβ40G33L, Aβ40G37L, and the C-terminal 20-residue fragment AV20. Structural insights into the interactions of these peptides with NDs, that serve as membrane mimics have been investigated using nuclear magnetic resonance (NMR) spectroscopy, fluorescence spectroscopy, molecular dynamics (MD) simulations, CD spectroscopy, etc. The elucidation of the GxxxG motif in Aβ aggregation kinetics, this research aims to advance our understanding of AD pathogenesis and potentially contribute to targeted therapeutic strategies.

**Figure 1.**
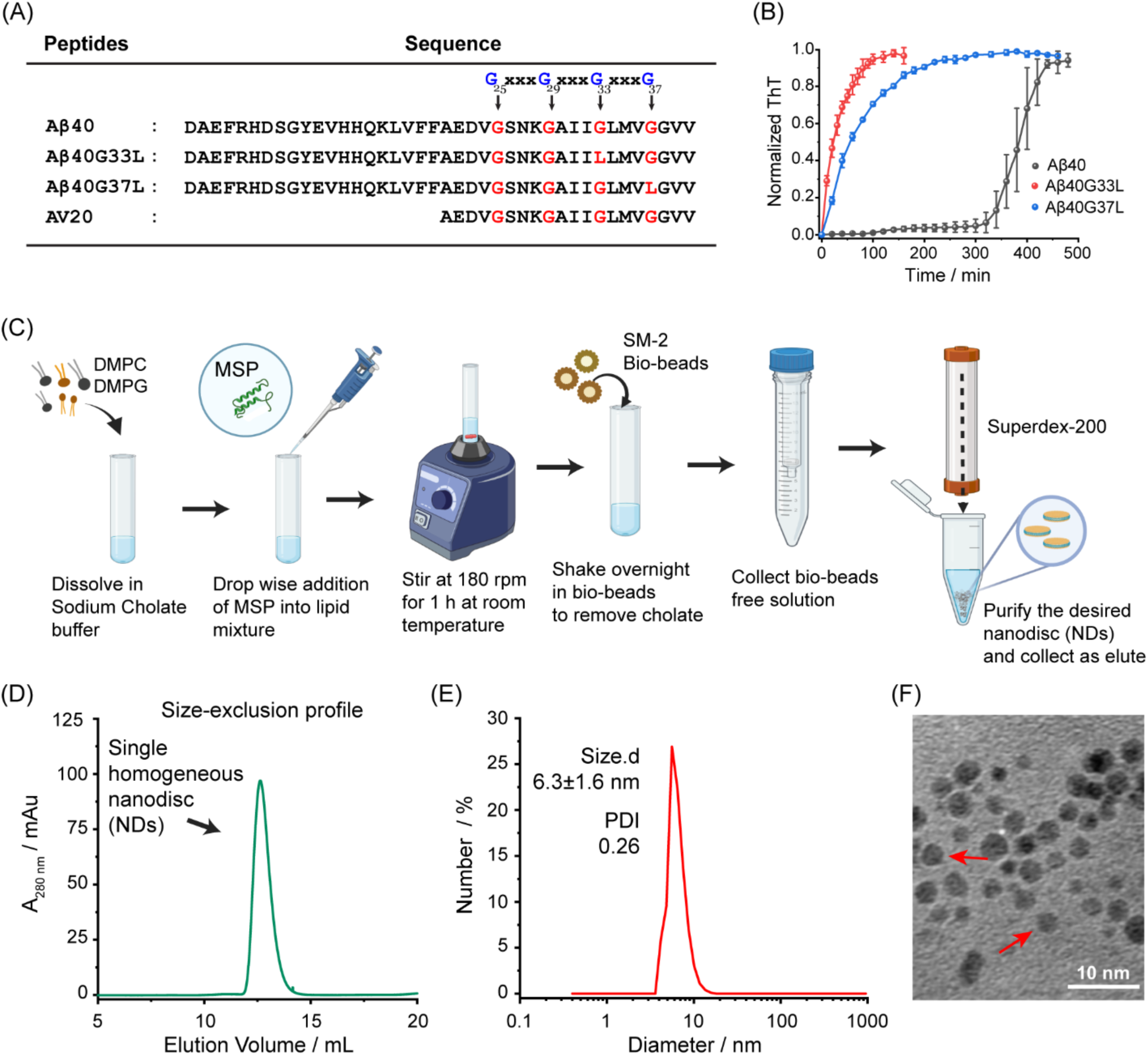
Set of peptides and characterization of the prepared nanodiscs. (A) Amino acid sequences of Aβ40 peptide, GxxxG motif mutants (Aβ40G33L, Aβ40G37L), and C-terminal fragment of Aβ40 (AV20). (B) Aggregation kinetics of Aβ40, Aβ40G33L, and Aβ40G37L investigated by ThT-based fluorescence assay at 37°C, 250 rpm sharking condition. Aβ40 displayed a sigmoidal growth kinetics with an observable lag-time, whereas the mutant peptides (Aβ40G33L and Aβ40G37L) had no lag-time and displayed rapid aggregation compared to the WT Aβ40. All experiments were repeated three times, and the data were averaged (±SD). (C) Schematic illustration of nanodisc (NDs) preparation generated using BioRender.com. NDs composed of DMPC and DMPG (3:1) was prepared using a MSPD1ΔH5 variant to wrap around the assembled lipid molecules. (D) Single, narrow and symmetric well-defined peak at Size Exclusion Chromatography (SEC) profile confirms the formation of homogeneous NDs. (E) Dynamic light scattering (DLS) of the NDs showed the hydrodynamic diameter of the formed NDs to be around 5-8 nm. (F) TEM image confirms the uniformity of shape and size of the purified NDs.

## 2. RESULTS AND DISCUSSIONS

### 2.1. GxxxG Motif Regulates Aβ40 Self-Assembly Kinetics

We examined the aggregation kinetics of monomeric Aβ40 (Figure S1 and S2) as well as the effect of Gly→Leu mutations at positions 33 and 37 within the GxxxG motifs on the aggregation kinetics of Aβ40 by using Thioflavin T (ThT) fluorescence assay (Figure 1B). While the ThT fibrillation kinetics for Aβ40 were in alignment with the sigmoidal growth model, consistent with previous findings reported by various research groups (45), (46)the mutant peptides behaved differently. On incubation at 37 °C, and 250 rpm, the saturation of Aβ40 aggregation was reached at approximately 420 min of incubation, with an observable lag time (t_lag_) of 356 ± 30 min and half-life (t_½_) of 380 ± 27 min. Previously Hung et al. have reported that the substitution of Gly→Leu at positions G33 and G37 resulted in a remarkable increase in the fibrillation rate of Aβ42 peptides (47). We used similar mutants for the Aβ40 peptides (Aβ40G33L and Aβ40G37L), and observed that in a similar experimental condition the mutant peptides exhibited rapid aggregation, achieving saturation within only 120 minutes of incubation (Figure 1B) without any observable aggregation t_lag_, and they did not follow sigmoidal growth curve, indicating their propensity for rapid nucleation and aggregation (Figure S3). The observed differences in aggregation behavior between the wild-type (48) Aβ40 and the mutants (Aβ40G33L and Aβ40G37L) suggested that these Gly residues play a crucial role in influencing the structural transitions as well as fibrillation dynamics of the Aβ40 peptides. Time-dependent circular dichroism (CD) spectroscopy showed the changes in the secondary conformation of the WT Aβ40 and mutant peptides during aggregation (Figure S4). Initially, Aβ40 displayed a random coil conformation (negative minima around 200 nm), which gradually transitioned into a β-sheet conformation (negative minima around 216 nm) as the aggregation reached saturation(49). In contrast, the mutant peptides displayed a remarkably swift structural transition, with a complete absence of any random coil structure even in the early-stages. In fact, within just 180 minutes of incubation, the mutants adopted into β-sheet conformation completely. Hence, both ThT assay and CD spectroscopy consistently demonstrate that the presence of Gly residues at positions 33 and 37 significantly impacts Aβ40 aggregation kinetics and its structural interconversion.

### 2.2. Preparation and Characterization of MSP-Nanodiscs

To investigate the role of the G^33^xxxG^37^ motif in membrane-mediated oligomerization of Aβ40, we prepared nanodiscs (NDs), disc-shaped lipid bilayers encircled by a membrane-scaffold protein (MSP) belt, representing near-native membrane mimetics, with tunable size and charge distribution. MSP-NDs have been reported to be significant in the study of membrane proteins and amyloid oligomeric intermediates (50) (51) (52). Several truncated variants of MSP are used to form nanodiscs of different sizes (50), (53). The size of the MSP has been found to play crucial role for long-term stability of the associated NDs as well as for obtaining high-resolution structural information of oligomers in solution NMR (50) (24).

In our study, we used the MSPD1ΔH5 variant, consisting of 167 amino acids (19.45 kDa), generating NDs in the size range of ∼5-8 nm (24) (Figure S5). The NDs were composed of DMPC (1,2-dimyristoyl-sn-glycero-3-phosphocholine) and DMPG 1,2-Dimyristoyl-sn-glycero-3-phospho-rac-(1-glycerol), at a 3:1 ratio (Figure 1C and Table S1), a lipid composition widely used to study peptide aggregation, including Aβ (28). In size exclusion chromatography (SEC), a single significant peak was observed at elution of 12.1-14.2 ml (Figure 1D) denoting the formation of homogeneous NDs (54). Dynamic light scattering (DLS) experiments (Figure 1E) and transmission electron microscopy images (Figure 1F) confirm the formation of uniformly sized MSP-lipid NDs within the size of 4-7 nm (Figure 1E and Figures S6A-B). One-dimensional ^1^H NMR of the MSP-lipid NDs showed no detectable peaks in the amide region (6.0 - 8.5 ppm), indicating the absence of free MSP protein in the solution (Figure S6C). This finding confirmed that all the MSP proteins successfully wrapped around the lipid bilayers, which is a characteristic feature of a well assembled MSP-lipid ND(50).

### 2.3. GxxxG-Mediated Membrane Interaction Modulates Aβ Aggregation Kinetics

To gain insights into the effects of NDs on Aβ40 aggregation kinetics and to elucidate the distinct roles played by G33/G37 residues, we first employed the ThT fluorescence assay at different lipid-peptide ratios (L:P), accompanied by coarse-grained (CG) MD simulations. Our findings revealed that both the lipid concentration and the Gly residues play crucial roles in regulating Aβ40 fibrillation. The effects of NDs on Aβ aggregation kinetics were found to be complex (Figure 2A), with a distinct pattern. At an equimolar ratio of lipid and protein (L:P=1), NDs promoted Aβ40 aggregation with a slight decrease in the t_lag_ (Figure S7). However, the decrease was not very significant. At higher lipid concentrations (L:P=10), the aggregation was enhanced in the presence of NDs. Intriguingly, with a further increase in lipid concentration (L:P=20), a substantial delay in Aβ40 aggregation was observed along with a significant decrease in the t_lag_ (Figures 2A and S7). Subsequent increments in the lipid concentration (L/P=50 and 100) resulted in almost no increase in the ThT signal, indicating the absence of Aβ40 fibrillation in the presence of high concentrations of NDs (Figures 2A and S8). No increase in the ThT intensity was observed for NDs alone (Figure S8).

**Figure 2.**
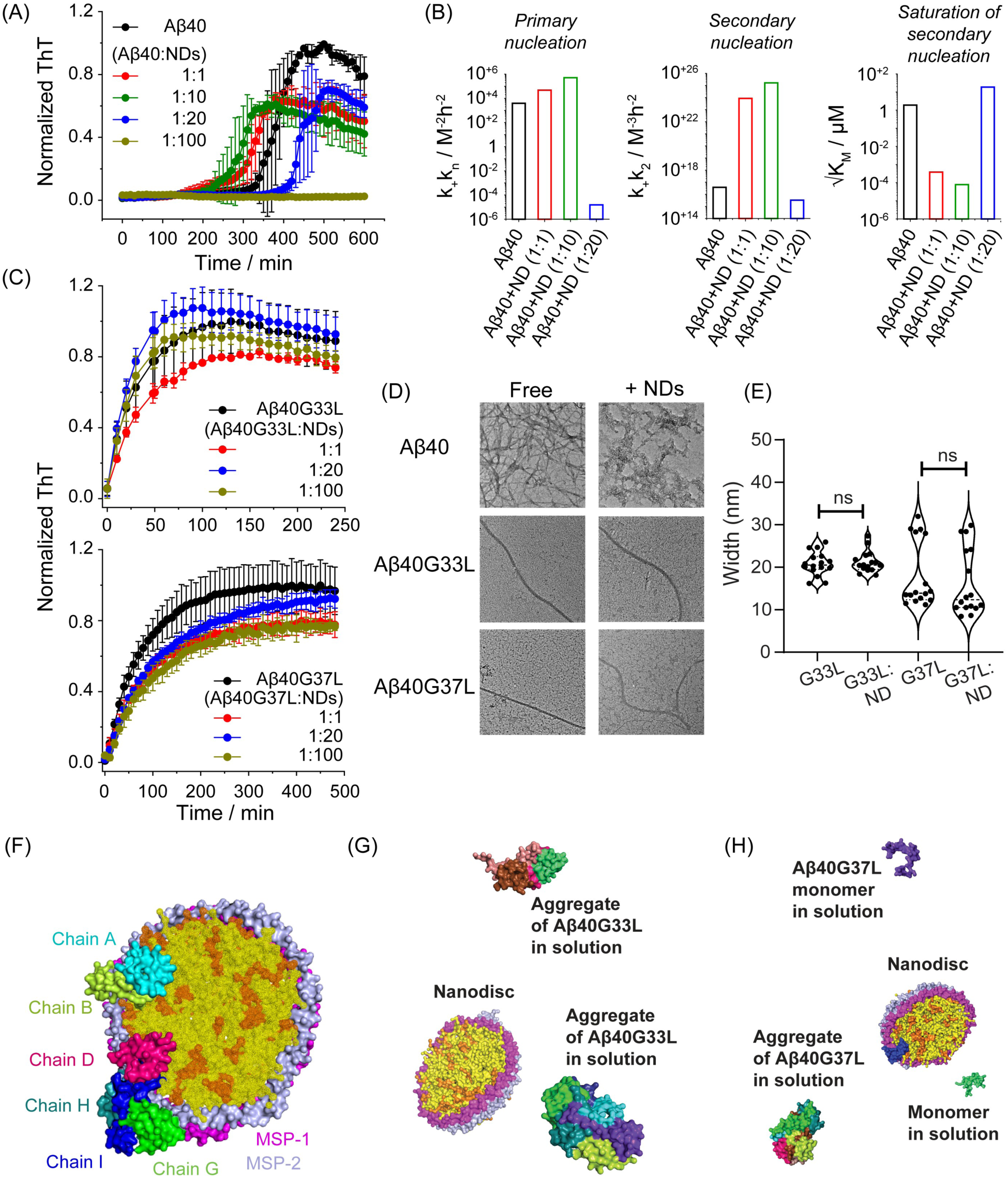
GxxxG motif modulates membrane-associated amyloid aggregation. Aggregation kinetics of (A) Aβ40 (20 μM) in the absence and presence of different lipid ratios investigated by ThT fluorescence assay at 37 °C under shaking (300 rpm) conditions in a plate reader. Presence of NDs displayed a complex kinetic behavior on the aggregation of Aβ40 under different lipid ratios. All experiments were repeated three times, and the data were averaged (±SD). (B) Global fitting of the aggregation kinetics data of Aβ40 in the absence and presence of different lipid ratios using multi-step secondary nucleation model. From the fitting of the data, parameters like the combined rate constants (k_+_k_n_ and k_+_k_2_) and saturation of secondary nucleation (√K_M_) were obtained and plotted in a bar diagram. (C) Aggregation kinetics of two mutants Aβ40G33L (20 μM), and Aβ40G37L (20 μM) in the absence and presence of different lipid ratios investigated by ThT assay at 37 °C under shaking (300 rpm) conditions, showing no specific effect of NDs on the aggregation kinetics. All experiments were repeated three times, and the data were averaged (±SD). (D) TEM images of Aβ40, Aβ40G33L, and Aβ40G37L in the absence and in presence of NDs (L:P=20), displaying the significance of G33 and G37 residues in membrane interaction and differential aggregation. (E) Analysis of the widths of Aβ40 mutant fibrils (G33L and G37L) in the absence and presence of NDs using ImageJ software. No significant morphological change was observed in the presence of NDs. Significant changes are marked with *(p ≤ 0.05) according to an unpaired t-test, while “ns” indicates non-significance. (F) Interaction of Aβ40 oligomers on NDs obtained from coarse-grained (CG) MD simulation. Aggregate formation of (G) Aβ40G33L, (H) Aβ40G37L in solution was observed without the formation of aggregate on NDs.

We further executed the fitting of the kinetic curves using the global fitting platform Amylofit(55), to compare the experimental kinetics data with the integrated rate laws, and obtained the rate constants associated with individual microscopic steps (Figure 2B). We used three independent curves for fitting each dataset using several models and selected the multi-step secondary nucleation model for our study as the observed residual mean error was the least (Figure S9)(45). We calculated the rate constants such as k_+_ (elongation rate constant), k_n_ (primary nucleation rate constant), k_2_ (secondary nucleation rate constant), as well as the Michaelis constant K_M_ (Figure 2B, Figure S10). At lower lipid concentrations, there was a significant increase in both the combined rate constants for primary (k_+_k_n_) as well as secondary (k_+_k_2_) nucleation. On the contrary, when the lipid concentration was higher, the rate constants decreased significantly, which directly reflected in the aggregation kinetics (t_lag_ and t_half_) (Figure S10). The observed values of k_+_k_2_ were higher than k_+_k_n_ (k_2_/k_n_ > 10^12^). The ratio of k_2_/k_n_ consistently increased with the increase in the lipid concentration, suggesting that most of the new aggregates were formed through surface-catalysed secondary nucleation rather than primary nucleation. The effects of NDs on Aβ aggregation kinetics exhibited a classic concentration-dependent biphasic pattern. At lower lipid ratios (L:P=1 and L:P=10), the NDs acted as catalytic surfaces, locally concentrating the Aβ monomers and significantly promoting secondary nucleation. Conversely, at higher lipid concentrations (L:P ≥ 20), we observed a drastic increase in the lag time. This inhibition is attributed to a surface dilution effect; the vast excess of available membrane surface isolates the peptides across multiple nanodiscs, reducing the probability of peptide-peptide collisions required for fibril elongation.

On the other hand, NDs did not have any distinct impact on the aggregation kinetics of mutant peptides (Aβ40G33L and Aβ40G37L) (Figure 2C) even at higher (L/P) concentrations. These findings highlight that the Gly residues (G33 and G37) significantly modulate the Aβ40 fibrillation. Further, there is an intricate interplay between the GxxxG-mediated membrane interactions and Aβ40 aggregation kinetics.

To visualize the effect of NDs on fibril formation by Aβ40 and its mutants, we performed transmission electron(16) microscopy imaging of Aβ40 and the mutants in both the absence and presence of NDs (Figure 2D). Aβ40 exhibited a dense network of amyloid fibrils in the absence of NDs of width ∼23 nm ± 4.3 nm (Figure S11). In contrast, in the presence of NDs, Aβ40 did not form complete fibrils and instead remained as disordered aggregates (Figure S12). The mutant peptides (Aβ40G33L and Aβ40G37L), on the other hand, existed as long isolated fibrils both in the absence and presence of NDs of width ranging from 21 nm ± 3 nm for Aβ40G33L and 18 nm ± 7 nm for Aβ40G37L (Figures S13–S17). The overall width of the mutant fibrils remained unchanged in the presence of NDs (Figure 2E).

The aggregation patterns of wild-type Aβ40, Aβ40G33L, and Aβ40G37L were also studied in detail using coarse-grained (CG) MD simulations, each performed for a 2 μs timescale. Two MSP molecules (MSP-1 and MSP-2), with a diameter of ∼98 Å, together with DLPC and DLPG lipid molecules (3:1), were used to prepare the NDs. Ten Aβ40 monomers (chains A–J) were used in the model system (Figure S18A). During the simulation, Aβ40 monomers formed self-aggregates in solution. These aggregates further interacted with the lipid moieties and MSP molecules of the NDs during the MD simulation timescale (Figure 2F). While some Aβ40 aggregates were associated with NDs, soluble aggregates and single monomers were also present in solution (Movie S1). We also investigated the roles of G33 and G37 using MD simulation. Both Gly residues were mutated to Leu, and each mutant was simulated separately using a model similar to that used for WT Aβ40.

In contrast to Aβ40 peptides, the mutant variants (Aβ40G33L and Aβ40G37L) formed only self-aggregates in solution, and these aggregates did not attach to the NDs (Figures 2G-H). These behaviors were observed in triplicate for both mutants (Figures S18B–E). Overall, our findings show that NDs influence the aggregation propensity of Aβ40, whereas in the mutants, NDs play a negligible role.

### 2.4. Residue Specific Interaction Between Aβ40 and Nanodiscs by Solution-NMR

To obtain an atomistic-level insight into the binding mechanism of Aβ40 with NDs, we used a combination of 1D, 2D, and 3D NMR experiments in the solution-state. One-dimensional ^1^H NMR spectra showed that upon addition of NDs (L:P=20), the amide proton intensity decreased instantaneously by ∼10 % (Figure S19). The NMR spectra further revealed a time-dependent decrease in amide proton signal intensity, reaching approximately 35 % after 44 h. Remarkably, significant CSPs were observed after this time point, while the overall signal intensity remained almost unchanged.

Next, we performed the 2D ^1^H-^15^N heteronuclear single quantum coherence (HSQC) NMR of Aβ40 in the presence of NDs at 25 °C and the spectra were assigned based on our previous study(56) (38). On interaction with NDs (L/P=20), Aβ40 initially showed a minor decrease in the peak intensity without any notable CSPs (Figure S20). With time, however, significant decay (25 %-35 %) in the NMR peak intensity was observed for residues, mainly encompassing the central hydrophobic and C-terminal regions (Figure 3A-B). A similar effect was previously reported by our group,(38) in which preformed fibrils seeded Aβ40 aggregation through reversible binding to the fibril surface, thereby directly modulating the exchange rates between the unbound and bound states on the NMR timescale. In this case also, the drop in the signal intensity was uneven throughout the peptide backbone, and hence it could be inferred that the signal decrease was not solely due to the formation of amyloid fibers. This drop in the intensity of the amide proton signal can be attributed to the intermediate exchange between the unbound and bound forms in the NMR timescale. Following 44 h of incubation, the overall signal intensity remained constant; however, significant CSPs were observed from this point, suggesting a fast exchange between the bound and unbound forms in the NMR timescale that results in a shift in the peak position along with the occurrence of oligomers with alternative conformations (Figures 3A-C, S21) (38). These oligomeric conformers remained stable for a long period of time before there was a complete broadening of the peak intensity.

**Figure 3.**
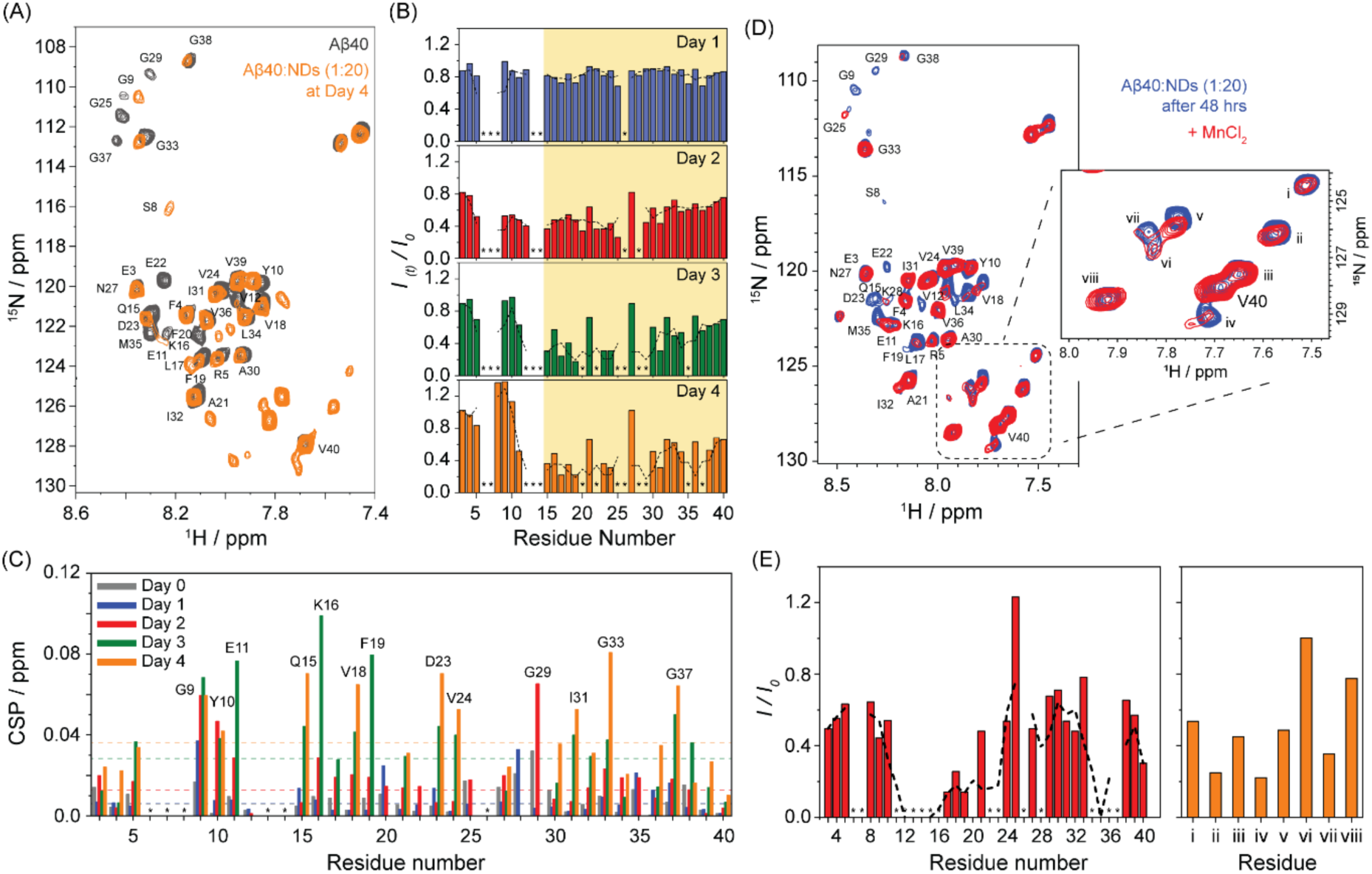
Interaction between Aβ40 and NDs by solution NMR. (A) Overlay of 2D ^1^H-^15^N HSQC spectra of a freshly prepared Aβ40 (∼80 µM) sample (black) and with nanodiscs (L/P=20) after formation of oligomers (orange) at 25 °C. Significant CSP and as well as new conformational peaks were observed in the vicinity of V40 in the presence of nanodiscs (L/P=20) after 48 hours of incubation. (B) Time-dependent changes in the peak intensities along the peptide backbone measured from 2D ^1^H-^15^N HSQC spectra of Aβ40 (∼80 µM) sample in presence of NDs (L/P=20) at 25 °C. Points represent the signal intensity of the peak maxima (*I(t)*) relative to the signal intensity in the absence of NDs (*I0*). Black dashed line represents the running average. Central hydrophobic region and C-terminal region showed maximum signal intensity drop. (C) Chemical shift perturbations (CSPs) of Aβ40 over time following the addition of NDs (L/P=20). The CSP was taken as the difference between the time point before and after NDs addition. Dashed horizontal lines are the average CSPs. (D) 2D ^1^H-^15^N HSQC spectra of Aβ40 in the presence of NDs (blue) after 48 h of incubation at 25 °C, and followed by titration with the paramagnetic quencher, MnCl_2(43)_. (E) Relative signal intensities of Aβ40 residues were measured in presence of MnCl_2_ (0.5 mM) for quantifying solvent accessibility of ND-bound Aβ40. N-terminal, C-terminal residues along with the newly formed peaks were partially embedded in the lipid bilayer The dashed line in the plots represents the running-average. The symbols (*) denote residues whose resonances did not display discernible resonances. All NMR experiments were done in Bruker Avance III 700 MHz spectrometer equipped with a 5 mm RT probe.

In the case of ND bound Aβ40, new peaks were detected in the 2D HSQC spectrum in the vicinity of V40, suggesting the generation of new oligomeric species on the surface of the NDs. This experiment was repeated more than four times, and with different batches of peptide. Although not identical, significantly close resemblance was observed (especially the formation of new peaks in the vicinity of V40), suggesting that these oligomers are structurally similar in nature (Figure S22). However, differences in the chemical shift between these different sets also indicates a complex oligomerization process where oligomers might be in a dynamic equilibrium.

Next, paramagnetic relaxation enhancement(57) NMR was conducted on these oligomers to identify the residues involved in membrane interaction. Mn^2+^ ions can selectively quench the NMR signals of solvent-exposed residues by enhancing their respective T_2_ relaxation rates, while residues embedded within the membrane bilayers are shielded from Mn^2+^ quenching. PRE experiment in the presence of 0.5 mM MnCl_2_ revealed that both the N-terminal and the C-terminal residues are partially embedded in the lipid bilayer (Figures 3D-E). The proton signal intensity of the amino acids spanning the central hydrophobic region was quenched, indicating that these amino acids are exposed to the solvent and dynamically interact with the surface of the NDs as well as the unbound form of the peptide. Additionally, the newly formed peaks were found to be partially protected from the quencher, suggesting that these alternative conformers were involved in active interaction with the lipid bilayer. The aggregation of Aβ40 in the presence of high lipid concentrations (L/P= 50) also showed a delay in NMR spectroscopy (Figures S23-24). Similar new peaks were detected in the HSQC spectra in the vicinity of V40, but at a later time point compared to free Aβ40, further confirming ND-mediated distinct oligomer formation.

### 2.5. Atomistic Details of Aβ40-NDs Interaction Using MD Simulations

To elucidate the detailed structural basis of the interaction between Aβ40 and MSP-based ND, an all-atom MD simulation was performed, containing ten Aβ40 monomers (labelled as chain A to chain J) in the model system (Figure 4A) like the CG simulation discussed earlier. Though the 150 ns all-atom simulation process is not sufficient to understand the complete Aβ40 aggregation process on ND, initial attachment and interactions of Aβ40 oligomer and monomer with MSP and lipid molecules were observed in the timescale. Among the ten Aβ40 monomers (chain A to J) used in the simulation, two of them (chain I and chain J) remained in the solution till 150 ns (Figure S25), seven monomers were present in an aggregated form associated with the ND, and chain E interacted with MSP (Figure 4B). Interestingly, V24-N27 of the C-terminal region of Aβ40 was observed to play a major role in the attachment and stabilization of the aggregate on ND (Figure 4C). Additionally, S26 and N27 formed hydrogen bonds with the phosphate group of the DMPC lipid molecules. In contrast, V24 formed a hydrophobic interaction with the methyl group of the lipid molecules. S8 and Y10 from the N-terminal region also formed hydrogen bonds and hydrophobic interactions with the ND, respectively (Figure 4C). The initiation and stabilization of Aβ40 aggregates in the solution were up to 120 ns. The closer proximity of the C-terminal region of chain A to the lipid molecules of the ND draws the Aβ40 aggregates to interact with the ND (Figure 4E, Movie S2). These Aβ40-ND interactions mediated by chain A remain stable, and the aggregate remains attached to the ND up to 150 ns (Figure 4E). Chain E interacts with the MSP proteins within 50 ns and remains attached throughout the simulation timescale (Figure 4F). At 150 ns, I31, I32, L17 and F20 of chain E were found to form hydrophobic interactions with L200 and L203 of MSP-1. A21 of the chain E was also part of the hydrophobic interaction with the methyl group of T197 of MSP-1. The E205 of MSP-1 formed a polar interaction with the side chain of H13 and the main chain of Y10, while K208 of MSP-1 formed hydrogen bonds with the side chain of S8. D1 of the N-terminal of chain E formed a salt bridge interaction with R215 of MSP-1, E3 formed a polar interaction with D1, although in a few frames, it interacts directly with the residues of MSP-2 (Figure 4D). During this simulation process, the C-terminal region of chain E also started forming a helical structure between K28-L34, besides maintaining the N-terminal helix. Both helices are present during the overall simulation timescale (Figure S26).

**Figure 4.**
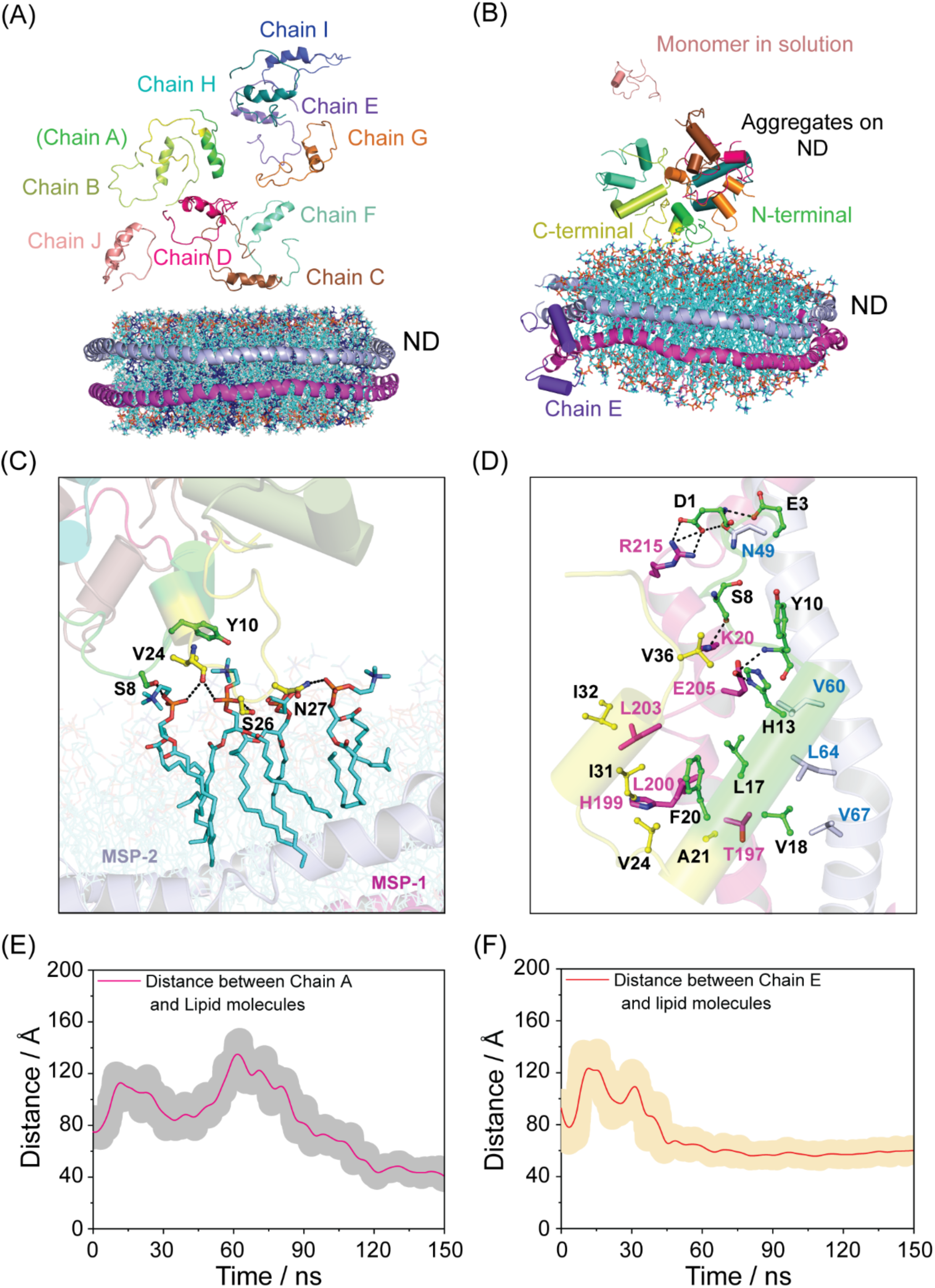
Interaction between Aβ40 and NDs using MD simulation. (A) Initial structure of ND containing MSP proteins and DMPC and DMPG lipid models and ten monomers of Aβ40 (chain A to J). The DMPC and DMPG molecules are shown in cyan and blue cartoon representation, respectively. The two MSP molecules (MSP-1 and MSP-2) are represented as magenta and light blue cartoon, respectively. All Aβ40 chains are color coded in cartoon representation. (B) MD simulation snapshot of Aβ40 aggregation on NDs. The Aβ40 monomers are shown in multiple colors in cartoon representation. Lipid molecules are shown in cyan line representation. (C) Interaction of chain A with DMPC and DMPG lipid molecules of ND observed during simulation process. The residues of Aβ40 and lipid molecules involved in interactions are shown in sticks. The N-terminal region of chain A is shown in green and C-terminal region is shown in yellow cartoon. (D) Interaction of chain E with MSP molecules of NDs. The residues involved in polar and hydrophobic interactions are shown in sticks. (E) Distance between center of mass (COM) of chain A with COM of lipid molecules during the all-atom simulation process. (F) Distance between COM of chain E with COM of lipid molecules during the all-atom simulation.

The residues of chain E, I and, J displayed the maximum fluctuations compared to the other monomers (Figure S27). This could be attributed to the differential preference of chain E in interacting with the MSP molecules over the lipid molecules of ND. Chain I and chain J remain in the solution and also do not participate in membrane-associated aggregation, resulting in the huge dynamics (Figures S28 and S29). It is important to note the temporal distinction between our computational and experimental models. The 150 ns all-atom MD simulation is designed to capture the rapid, initial docking events and early-stage stabilization of Aβ on the ND surface. These early hydrophobic anchor points (e.g., V24, N27) serve as the nucleation seeds that ultimately drive the slow, large-scale conformational transitions leading to the stable oligomeric species observed in our 48 h NMR experiments. While our all-atom MD simulations reveal direct hydrophobic interactions between Aβ40 (e.g., residues I31, I32, L17, F20) and the MSP belt, this dual binding to both lipid acyl chains and scaffold proteins accurately reflects the heterogeneous, crowded nature of native neuronal membranes, where Aβ simultaneously interacts with lipid rafts and membrane receptors.

### 2.6. Restriction of Residue Specific Backbone Mobility of Aβ40 upon Binding with Nanodiscs

Since the binding of Aβ40 to the membrane affects its backbone dynamics or motional rigidity, we measured the residue-specific transverse relaxation rate (R_2_)(58) of Aβ40 in the absence and presence of ND, using a lipid-to-protein ratio of 20 (L/P = 20). These measurements were performed at different time points e.g., at the beginning, after 24 h, and after 48 h of incubation. The deviation in R_2_ values between free and ND-bound state is represented as ΔR_2_. Interestingly, a positive magnitude in residue specific ΔR_2_ profile (Figures 5A-B) suggests the motional rigidity and henceforth the direct involvement of those residues in interaction with NDs. ΔR_2_ at different time points of incubation was also calculated to understand time dependent residue specific differential binding of Aβ40 to membrane surface. During the initial time point (up to 24 h), there was a negligible change in the transverse relaxation rates for some N-terminal residues (F4, R5), C-terminal residues (V36, V39, V40), and some central-region residues (Y10, V12, E22, V24) (Figure 5A). However, there was a significant gradual increase in ΔR_2_ values for residues including F19, A21, D23, N27, A30, I32, and G38. This suggested that these residues were directly involved in binding to the ND surface and in surface-mediated oligomerization during the early stages. All atom simulation data well corroborate with the involvement of N27 residue with lipid surface. At later time, after 48 h or at the time of new peak appearance, interesting dynamic behaviour was observed among all residues. Decrease in ΔR_2_ values were observed for C- and N-terminal residues like E3, V36, V39, V40 and A21, N27, I32 indicating no significant interacting role at later stage of incubation, whereas greater change in ΔR_2_ values were observed specifically for residues Y10, V12, F19, D23, V24, A30, G38 (Figure 5B). All atom simulation data equally confirm the hydrophobic interaction of Y10 and V24 with the lipid molecules. MD simulation in conjunction with NMR confirms that some N- and C-terminal residues play a role in the early stages of membrane interaction; the most consistent and significant interactions occur through residues F19, A21, D23, N27, A30, and G38. These residues are involved in both binding to the membrane lipid surface and membrane-assisted oligomerization. This observation again corroborates with our chemical shift perturbation (CSP) and PRE data. On the other hand, the apparent difference between the PRE shielding and the decreased R_2_ values at the N- and C-termini perfectly aligns with a ‘tethered’ binding model. At 48 h, the central hydrophobic regions are rigidly anchored to the NDs or oligomeric core (high R_2_), while the termini retain high segmental mobility (low R_2_). However, because these mobile termini are tethered to the membrane surface, they exist in a restricted spatial volume close to the bilayer, experiencing transient shielding from the bulk solvent as captured by our PRE measurements. Interestingly, it has been observed that ΔR_2_ value of Y10, V12 and V24 increased suddenly at later stage (24-48 h), showing that probably these two residues are involved in the binding scenario at later time and are associated with the new surface mediated oligomeric conformation as also observed from 3D NMR (Figure S30). To identify the new peaks appearing near residue V40 in the 2D HSQC spectrum during incubation of Aβ40 with 3:1 DMPC/DMPG at a lipid-to-peptide ratio of 20:1 (L/P=20), we performed triple resonance NMR experiments, specifically HNCA and CBCACONH, to sequentially assign the backbone resonances of these new signals. Unfortunately, due to the lack of well-resolved resonances in CBCACONH spectra, we were unable to assign all the new peaks. We could unambiguously assign only two new peaks as V12’ and F20’ (Figure S30). Our simulation data identified the involvement of F20 in hydrophobic interaction with MSP molecule (Figure 5C). So, positive ΔR_2_ resulted not only for peptide-membrane interaction but also for protein-protein interaction as well as conformational changes.

**Figure 5.**
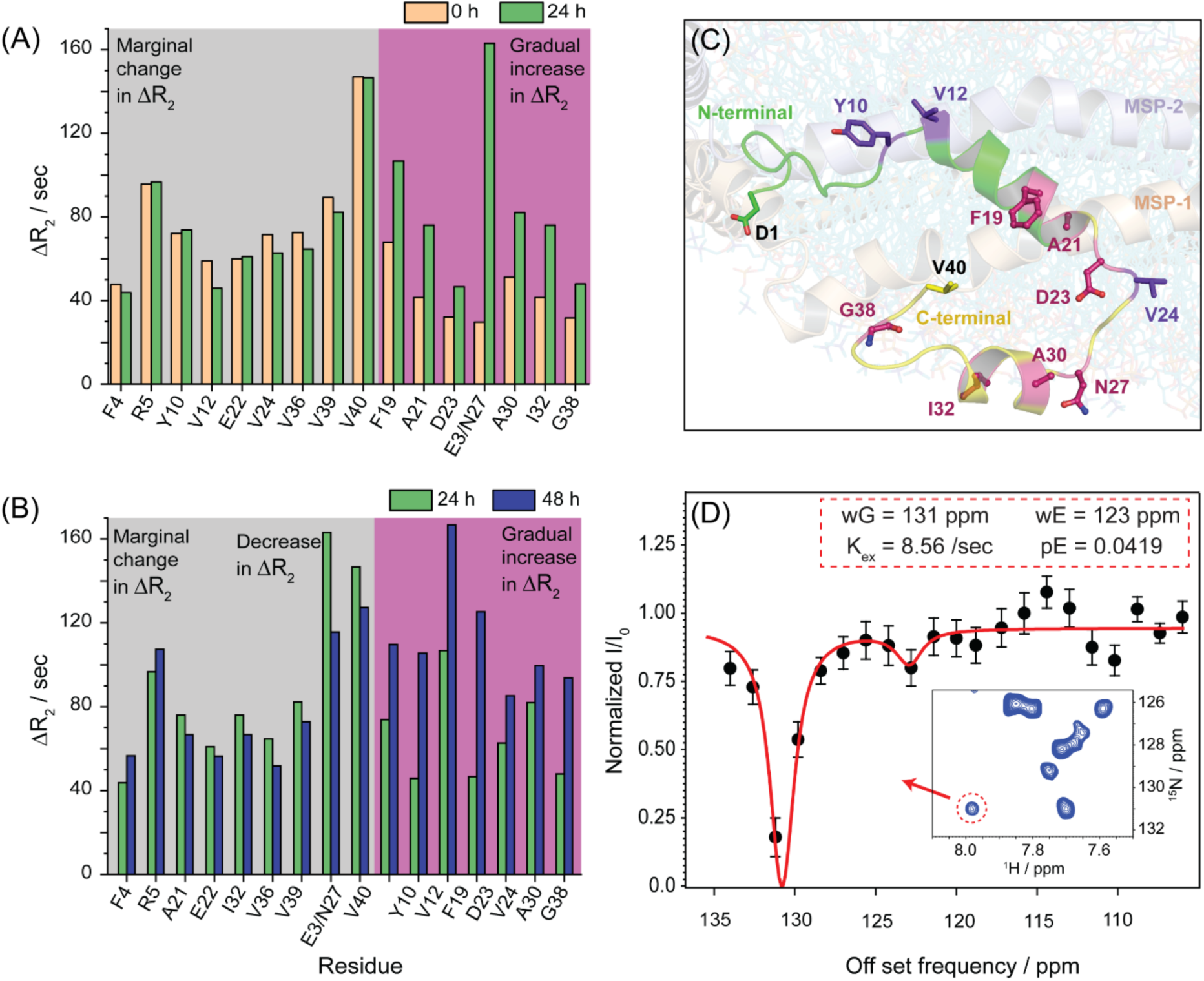
Restriction of residue specific backbone mobility of Aβ40 upon binding with NDs. The differences in transverse relaxation rates (ΔR_2_) of free Aβ40 and nano-disc bound Aβ40 (L/P=20) at different time points. (A) Residue specific ΔR_2_ change between 0 and 24 hrs. R_2_ values of 0 and 24 hours are marked in orange and green bar plot respectively (B) Residue specific ΔR_2_ change between 24 and 48 hrs. R_2_ values of 48 hours are marked in blue. Residues with marginal changes and gradual increase in R_2_ values are represented with grey and pink background respectively. Time dependent different R2 values of different residues indicate differential residual dynamicity of Aβ40 on lipid surface. (C) The residues with significant changes of ΔR_2_ are shown in sticks and the structure of Aβ40 are shown in cartoon representation. F19, A21, D23, N27, A30, I32, and G38 might be involved in constant interaction with lipid environment and Y10, V12 and V24 might be involved in the generation of surface assisted new conformational oligomeric peaks. (D) CEST profile of one residue that is marked in red circle, showing two dynamic population ground or major state at 130.79 ppm and excited or minor state population at 122.86 ppm. Relaxation NMR experiments were done in Bruker Avance III 500 MHz spectrometer equipped with a 5 mm RT probe. CEST NMR experiment was done in Bruker Avance III 700 MHz spectrometer equipped with a 5 mm RT probe.

Chemical Exchange Saturation Transfer (CEST) NMR has been widely employed to characterize low-population, sparsely populated excited states of proteins that are in dynamic equilibrium with their predominant ground state.^[45]^ Hence, we employed CEST NMR to investigate how the new conformational entity of Aβ40 is involved in chemical exchange with the original conformation. Considering the two-state mechanism, the CEST effect for one of the new peaks (denoted in red circle in the inset HSQC spectra of Figure 5D) was fitted for the I/I_0_ data at every offset ppm value using Bruker’s Dynamic Center 2.8.6. We obtained the major chemical shift resonance at 130.79 ppm and its minor or excited population state resonating at 122.86 ppm. Of note, the ground state conformation was the major population (pG =0.9581) and the excited state conformation was the minor population (pE = 0.0419). Their chemical exchange rate, k_ex_, is 8.56 Hz (Figure S31), a slow exchange process,^[47]^ as 8.56 Hz < 70× (130.79 - 122.86) Hz. The resonance of the excited state (ω_E_ = 122.86 ppm) of the new peak (peak 1 in Figure S31A) closely matched the ground state resonance (ω_G_ = 121.996 ppm) of L34 (peak 16 in Figure S31A). This suggests that the new conformational peak (denoted in red circle in HSQC spectra of Figure 5D inset) is in chemical exchange with L34. Simulation data further support this, showing that L34 plays a significant role in forming the C-terminal helix upon interaction with the lipid membrane. The interaction with ND induces a widespread structural rearrangement, generating a distinct set of new NMR resonances that correspond to a stable, tethered oligomer. Through a combination of triple-resonance NMR, CEST, and intensity buildup tracking, we mapped multiple residues participating in this new conformation. Specifically, V12 and F20 were assigned directly, CEST revealed a slow-exchanging excited state for L34, and matched kinetic profiles confirmed the participation of G25, G33, and G37. Together, these map the core of the newly formed membrane-associated oligomer.

The attribution of the newly observed NMR peaks to surface-catalyzed oligomeric species, rather than alternative monomeric bound states or degradation artifacts, is supported by both exchange kinetics and sample stability. Equilibrium binding of monomeric Aβ to lipid surfaces typically occurs rapidly upon mixing, falling within fast or intermediate chemical exchange regimes. In contrast, the new resonances in our HSQC spectra emerge only after a prolonged 48 h incubation, demonstrating a time-dependent intensity buildup that inversely mirrors the decay of the unbound monomeric signal. Furthermore, CEST NMR analysis of the new conformation reveals a slow chemical exchange rate (*kex* = 8.56 Hz), pointing to a high activation energy barrier characteristic of stable higher-order oligomerization rather than transient monomeric docking. Additionally, the robust nature of the MSP ND scaffold, which maintains bilayer integrity for over 30 days, alongside standard validations of intact full-length Aβ40, definitively excludes ND disassembly or peptide proteolysis as the source of these spectral variations. Although oligomerization and binding to a massive ND complex typically restrict global molecular tumbling and induce uniform, severe line broadening, the surprisingly narrow line widths observed for specific residues (e.g., near V40) are fully consistent with a “tethered” or “fuzzy coat” binding model. In this structural paradigm, specific central hydrophobic segments (such as residues F19 to G38) act as rigid anchors that tightly interact with the ND and form the oligomeric core, leading to localized line broadening and elevated transverse relaxation (R_2_) rates. Meanwhile, regions outside these anchor points, particularly the extreme N-and C-termini, project outward into the bulk solvent and retain high segmental mobility. This rapid, localized spatial motion effectively averages out chemical shift anisotropy and dipolar couplings, allowing these specific unbound segments to produce sharp, narrow NMR peaks even while part of a massive, slowly tumbling oligomeric complex.

### 2.7. AV20-Membrane Interaction Resembles with Aβ40

AV20 (A21 – V40), the C-terminal peptide fragment of Aβ40, was previously found to be significantly toxic, in which the role of the GxxxG motifs were extensively studied(39) (43). Here, we observed the role of membrane interaction with AV20 in order to further explore the significance of the GxxxG motifs in oligomer formation. Both Aβ40 and AV20 interacted with the ND in a similar way. ThT fibrillation assay indicated that the peptide AV20 in the presence of NDs, rapidly underwent aggregate formation, which were comparatively stable (Figure 6A). The formation of AV20 aggregates and their interaction with nanodiscs were also observed through coarse-grained MD simulations over 2 μs. Similar to wild-type Aβ40, AV20 aggregates interacted with both MSP scaffold proteins and lipid molecules, as well as with each component individually (Figure 6B, Movie S3). To further characterize these interactions at atomic resolution, solution-state NMR studies were conducted to investigate the binding mechanism between AV20 and nanodiscs. While titrating with NDs, the peptide underwent broadening of the amide proton signal upon subsequent interaction (Figures S32 and S33). No immediate chemical shift perturbation (CSP) was visible. However, upon binding with NDs, AV20 showed significant CSPs over time and remained stable for a comparatively longer duration (Figure 6C). This can be attributed to the conformational transition and formation of stable oligomers. For further insight into the residue specific interaction between AV20 and nanodisc, 2D TOCSY NMR experiments were performed with AV20 in the presence and absence of NDs (Figure 6D). The residues spanning G^33^LMVG^37^ was found to be highly perturbed, suggesting significant involvement of this segment upon membrane interactions (Figure 6E). Apparently, the CSI plot indicated mostly random coil conformation, except for residues L34 and M35, which showed inclination towards helical conformation (Figure S34). Next, to get an exclusive insight into the Gly residues, we performed 2D HSQC for the Gly-labeled (^15^N labeled Gly residues) AV20 in the presence of NDs, at low temperature (10 °C) as a function of time. The interaction with nanodiscs induced distinct changes in the local magnetic environment for specific residues, reflected by upfield shifts for G25 and downfield shifts for G33 and G37. The emergence of these discrete new peaks, coupled with CEST NMR data, confirms the presence of an alternative, stable conformation in slow exchange with the unbound state (Figure 6F). The intensity-buildup curves for the three new peaks associated with residues G25, G33, and G37 closely resemble the decrease in intensity of the peaks corresponding to their unbound forms, indicating that these peaks are in slow exchange. These results are fully consistent with our earlier observations, in which real-time NMR experiments were used to monitor f⁵N-labeled Aβ40 in the presence of Aβ40 fibril seeds(38). As in those studies, the additional peaks are absent in samples incubated without fibers, supporting a mechanism of surface-mediated catalysis of Aβ40 aggregation. Brender et al. further demonstrated that these peaks do not arise from impurities or peptide degradation, but instead reflect the formation of new oligomeric species generated through interactions with fibrillar surfaces. Taken together, these findings indicate that, in this case, interaction with the ND surface promotes the formation of distinct oligomeric species, giving rise to the observed new peaks. Conversely, G29 and G38 exhibited only a decrease in intensity. As mentioned earlier, the emergence of new peaks could indicate a transitional state of monomers in a surface-catalyzed interaction leading to newer oligomeric species, (38) or it could reflect the binding of AV20 monomers to the NDs. Due to nanodisc-induced signal broadening, ab initio assignment of the new membrane-bound peaks is challenging. We therefore assigned the slow-exchange resonances to G25, G33, and G37 using three converging lines of evidence: (1) they emerged exclusively within the characteristic 15N glycine spectral region (108–114 ppm); (2) 7-day kinetic tracking (Figures 6F, G) revealed matching population dynamics, in which the exponential decay of the unbound peaks mirrored the intensity buildup of the new peaks; and (3) spectral analysis of Aβ40G33L and Aβ40G37L mutants retrospectively validated their identities. Collectively, these data confirm that these specific residues transition into a stable, tethered oligomeric state. Free AV20, on the other hand, showed a decrease in signal intensity, but no new peaks were observed (Figure S35).

**Figure 6.**
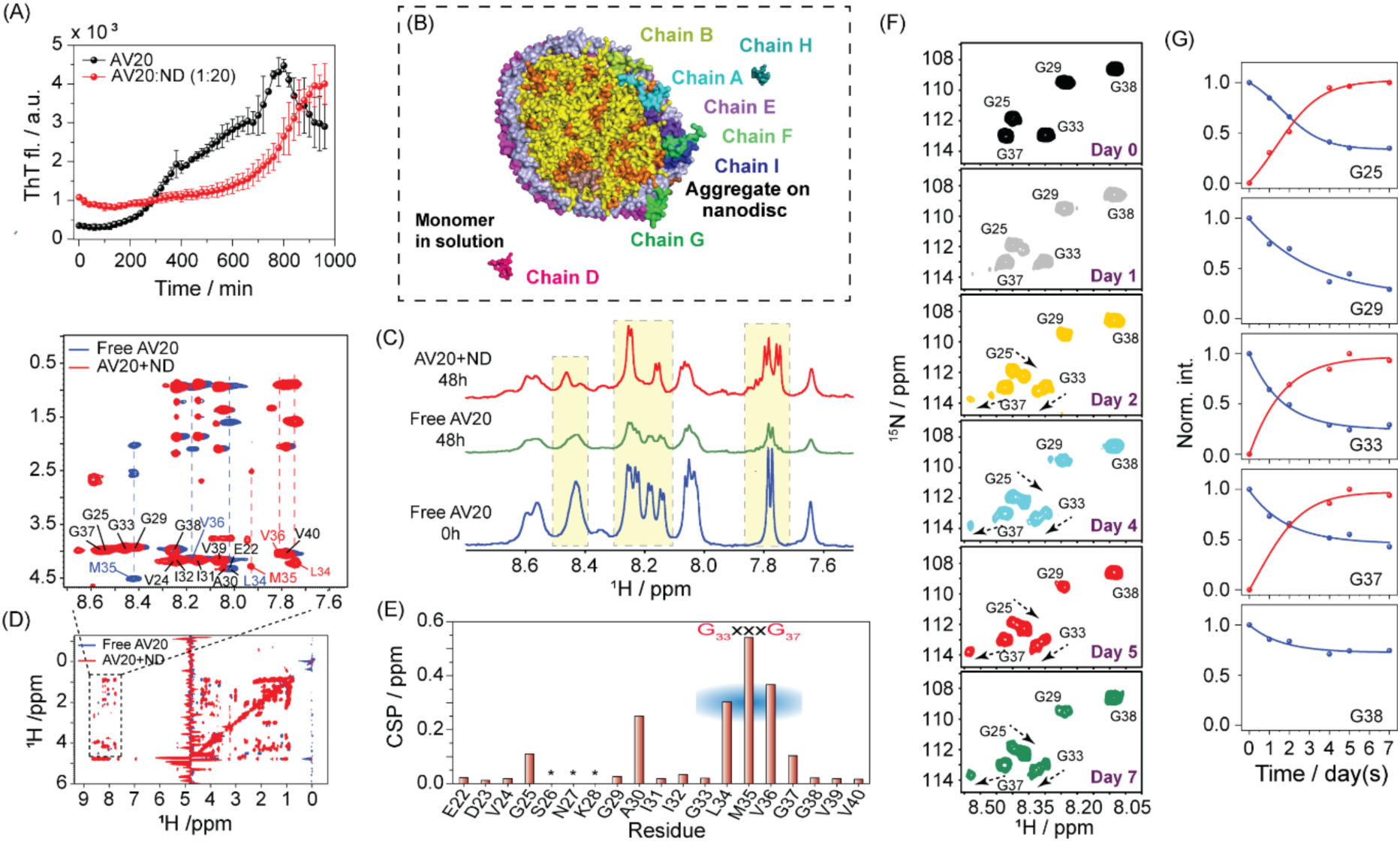
Structural insights of AV20-NDs interaction. (A) Aggregation kinetics of the C-terminal 20 residue Aβ40 fragment, AV20 (50 μM) in the absence and presence of NDs (L/P=20) by ThT fluorescence assay. All experiments were repeated three times at 37 °C under shaking (300 rpm) conditions in a plate reader, and the triplicates data were averaged (± SD). (B) Among ten AV20 monomers used in the CG MD simulation, eight of them forming aggregate on ND after 2 μs. (C) 1D (selected) ^1^H NMR spectra of free AV20 at 0 h, at 48 h and AV20 in presence of NDs at 48 h of incubation at 25 °C. While AV20 alone underwent line broadening, AV20 in presence of NDs showed both line broadening and significant CSPs. (D) 2D TOCSY NMR of AV20 in the absence (blue) and in presence of NDs (47), highlighting residue specific interactions. (E) The chemical shift deviation for Hα resonances (ΔHα) of each residue of AV20 in the presence of NDs from the free AV20. (F) Time-dependent 2D ^1^H-^15^N HSQC NMR spectra of AV20 in the presence of NDs (L/P = 20), focusing on the characteristic glycine region (108–114 ppm). Over the course of 7 days, the original unbound monomeric peaks gradually diminish, accompanied by the concurrent emergence of distinct, newly shifted peaks (indicated by dashed arrows) corresponding to the stable oligomeric conformation. (G) Normalized NMR peak-intensity trajectories as a function of incubation time. The exponential decay curves (blue lines) of the unbound G25, G33, and G37 resonances perfectly mirror the exponential intensity-buildup curves (red lines) of the newly formed slow-exchange peaks, illustrating a direct transition between the two structural states. All NMR experiments were done in Bruker Avance III 700 MHz spectrometer equipped with a 5 mm RT probe.

## 3. Conclusion

Our study provides a high-resolution mechanistic framework that clarifies the critical roles of G33 and G37 within the G^33^LMVG^37^ motif in regulating the aggregation kinetics of Aβ40. This study establishes that nanodiscs, which serve as near-native membrane mimetics, facilitate surface-catalyzed oligomerization of Aβ40 (Figure 7). Lipid concentration further modulates this process, with nanodiscs promoting the formation of relatively stable oligomers that are conformationally distinct from both monomeric Aβ40 and minor oligomeric species present in solution. Interactions between nanodiscs and both full-length Aβ40 and its C-terminal fragment (AV20) underscore the importance of the C-terminal region, particularly the G^33^LMVG^37^ motif. Structural analyses of Aβ40 and AV20 highlight the propensity of this domain to adopt helical conformations. Notably, mutations at G33 and G37 within the GxxxG motif disrupt these structural transitions, emphasizing their functional significance. It is important to note that oligomers formed in solution or *in vivo* may differ structurally and functionally from those generated in the presence of nanodiscs, and thus may not directly correspond to pathological species. Nevertheless, approaches such as the development of sequence- or conformation-specific antibodies, as well as small molecules that stabilize defined oligomeric intermediates, have proven valuable for therapeutic intervention. Overall, this work advances our understanding of membrane-mediated Aβ oligomerization and provides a foundation for the development of targeted therapies for Alzheimer’s disease and related disorders.

**Figure 7.**
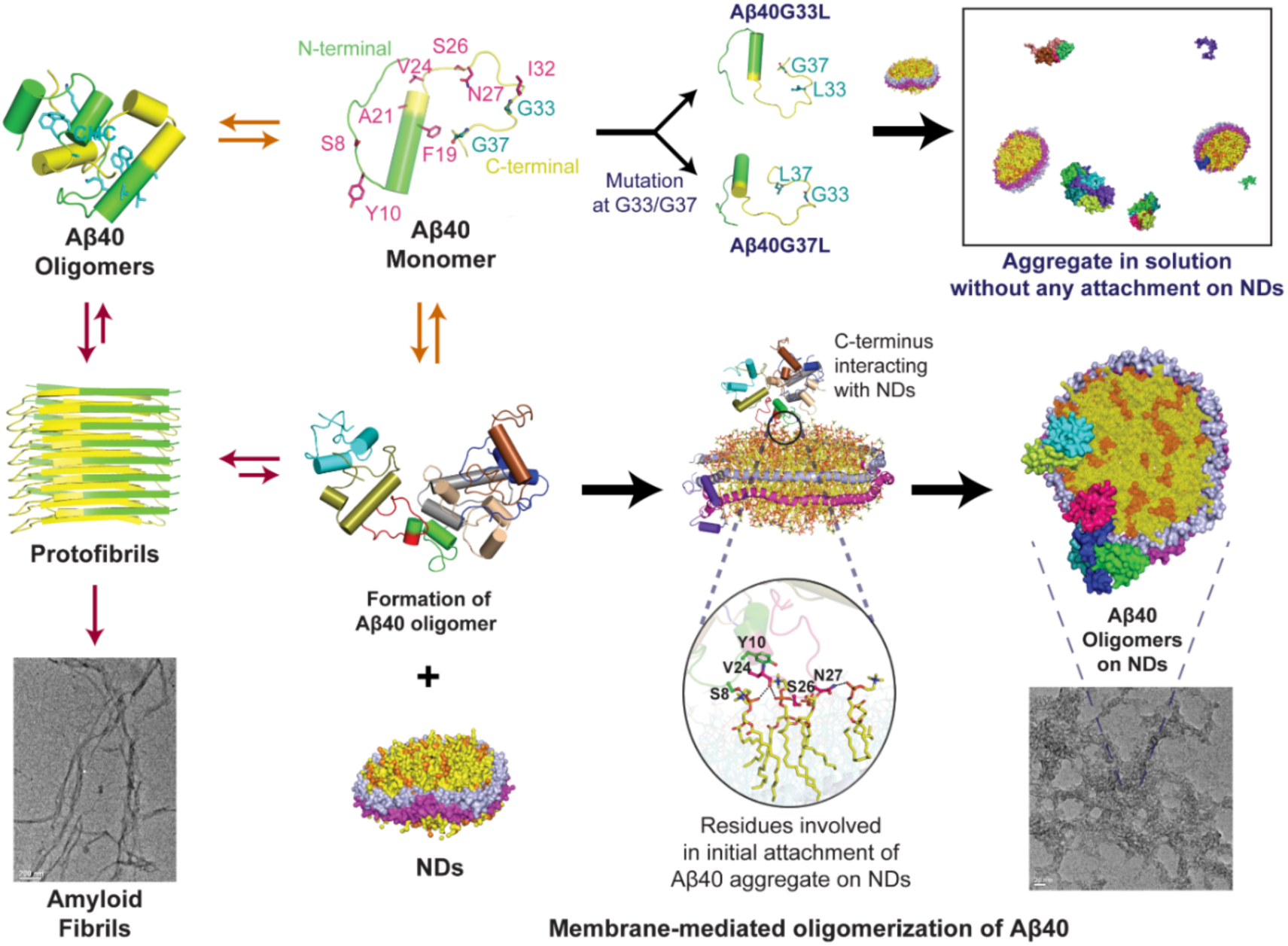
Schematic representation of membrane-mediated oligomerization of Aβ40. In aqueous solution monomeric Aβ40 self-assemble into oligomeric intermediates primarily via the central hydrophobic core (CHC). These oligomers are transient in nature and quickly convert into protofibrils and mature fibrils. In presence of NDs however, they form comparatively stable oligomers with distinct conformation. While a portion of Aβ40 self-assemble, the presence of NDs leads the C-terminal consisting of the GxxxG motif to interact with the lipid bilayer. This facilitates a conformational transition, where the N-terminal then insert into the membrane, resulting in a membrane-bound distinct oligomer. Mutation in the G33 and G37, on the other hand, hampers this interaction and leads to a faster aggregation into long isolated fibrils.

## 4. Experimental procedures

### 4.1. Preparation of Aβ40 and mutant peptides

Aβ40, Aβ40G33L, Aβ40G37L, and AV20 peptides were purchased from GenScript Inc. USA. The peptides were weighed and treated with 1,1,1,3,3,3-hexafluoro-2-propanol (HFIP) (Sigma-Aldrich, USA) or NaOH to obtain monomeric forms of the peptides. The treated samples were then subjected to gentle vortexing, followed by sonication for 10 s. Subsequent aliquots were prepared, which were lyophilized for 48 h and stored at -20 °C for long-term use.

### 4.2. Thioflavin T (ThT) fluorescence assay

Thioflavin-T fluorescence assays in 96-well black flat bottom plate using BMG LABTECH POLARstar Omega spectrometer (BMG Labtech, Ortenberg, Germany) or in cuvette using Hitachi F-7000 FL spectrophotometer (Tokyo, Japan).

Different conditions were used to prepare monomeric Aβ40, Aβ40G33L, and Aβ40G37L samples as mentioned previously. The aliquots were then thawed, followed by MALDI analysis and Size Exclusion Chromatography (SEC) to remove any fibrils that could seed the aggregation kinetics. The SEC-FPLC experiment was performed using a Sepharose 6 column (1 0mm internal diameter X 300 mm length) (Sigma-Aldrich) connected to an ÄKTA Pure FPLC system (GE Healthcare Life Sciences). The experiment was carried out at room temperature. The protein sample was dissolved in 500 µL of 20 mM Phosphate buffer (pH 7.4) containing 50 mM NaCl to a final concentration of 90 µM. The same buffer was used to equilibrate the column and as the running buffer for protein elution. Elution was carried out at a constant flow rate of 400 µL/min.

Stock solutions of Aβ40, Aβ40G33L, and Aβ40G37L were prepared by dissolving the peptides in a buffer (50 mM NaCl, 20 mM sodium-phosphate, pH 7.4), vortexing and ultra-sonicating, to obtain a concentration of 80 µM. To standardize the experimental conditions, the samples were incubated at 37 °C under both non-shaking as well as shaking conditions (100 rpm, 250 rpm, and 300 rpm). At different time intervals, different concentrations (2-20 µM) of aliquots were taken, mixed with ThT at a ratio of 1:1, and measured using BMG LABTECH POLARstar Omega spectrometer in a 96-well black flat bottom plate under shaking conditions (0 s before each 120 s cycle) at 37 °C. After standardization, the experiments were performed using 20 µM of the peptides, incubated with 20 µM ThT at 37 °C at 300 rpm.

The aggregation kinetics of the Aβ40 and mutant peptides were also studied using BMG LABTECH POLARstar Omega spectrometer in a 96-well black flat bottom plate. Equimolar concentration of ThT was added to the peptide sample and the fluorescence was monitored.

Three independent experiments were performed. The time-dependent ThT fluorescence data was fitted to the following sigmoidal growth model:

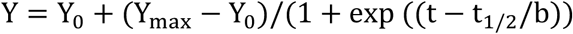

where t_1/2_ denotes the half-time, the time required to reach half of the fluorescence intensity, b is the apparent first-order constant and Y_max_, and Y_0_ are the maximum and initial fluorescence values, respectively. t_lag_ is the lag-time of the amyloid kinetics, and was denoted as (t_1/2_-2b).

### 4.3. Membrane scaffold protein (MSP) purification and Nanodisc preparation

To obtain a homogenous solution of uniformly-sized nanodiscs (ND), certain variables were optimized, which include size of membrane scaffold protein (MSP), amount of MSP, molar ratio of lipid/ MSP, surface area of each lipid molecules, and amount of lipids. For our study we have used the MSPD1ΔH5 truncate.

#### 4.3.1. Expression and purification of MSP

The expression and purification of MSP was performed using the protocol mentioned previously^[1]^. The pET28a-MSP plasmid (Addgene) was transformed into *E. coli* BL21 (DE3) cells. The protein was expressed in Luria Bertani (LB) medium, supplemented with 35 mg/L kanamycin. Protein production was induced by 1 mM Isopropyl-β-D-thiogalactopyranoside (IPTG) at 37 °C for 3-5 h. The harvested cell pellet was resuspended in MSP buffer (100 mM NaCl, 0.5 mM EDTA, 20 mM Tris-HCl, pH 7.5), centrifuged at 3,000 X *g* at 4 °C, and stored at -80 °C (50).

The cells were thawed, and resuspended in lysis buffer (500 mM NaCl, 1 % Triton X-100, 1 mM EDTA, 50 mM Tris-HCl, pH 8.0), sonicated and centrifuged. The supernatant was subjected to affinity purification on a nickel nitrilotriacetic acid resin (Ni-NTA) (Qiagen) column (Figure S2A). The column was washed twice with the same buffer, followed by washing with buffer containing 50 mM cholate. The protein was eluted in elution buffer (500 mM Imidazole, 500 mM NaCl, 50 mM Tris-HCl, pH 8.0) and dialyzed overnight at 4 °C against dialysis buffer (20 mM NaCl, 0.5 mM EDTA, 50 mM Tris-HCl, pH 8.0) in a dialysis tubing (10 kDa cutoff) (ThermoFisher Sc., Germany) (^53^). The purified protein was co-incubated with TEV protease and 1 mM Dithiothreitol (DTT) at 4 °C for 12 h to cleave the His-tag. After removal of the His-tag, the protein was dialyzed overnight, against a buffer containing 500 mM NaCl, 20 mM Tris-HCl, pH 8.0 and subjected to the Ni-NTA column. The flow through was collected and dialyzed against a buffer containing 100 mM NaCl, 20 mM Tris-HCl, pH 7.4. The protein was then concentrated using a 3.5 kDa membrane cutoff. Protein purity was checked using SDS Page and stored at -80 °C.

#### 4.3.2. Assembly of the Nanodiscs

3.22 mg of 1,2-dimyristoyl-sn-glycero-3-phosphocholine (DMPC) and 1.1 mg of 1,2-dimyristoyl-sn-glycero-3-phospho-(1’-rac-glycerol) were weighed and dissolved in a buffer (100 mM NaCl, 0.5 mM EDTA, 20 mM cholate, Tris-HCl, pH 7.5). Purified MSP was added dropwise to the lipid combination, with continuous stirring, and co-incubated at room temperature for 1 h (Figure 1C). The assembled mixture (Table S1) was then incubated in the presence of bio-beads, overnight, at room temperature, to remove any residual detergent. Next day, the bio-beads were centrifuged at 1,000 X *g*, and the mixture containing the ND was collected, which was subjected to further purification using size exclusion chromatography. The Superdex 200 10/300 increase GL column (GE Healthcare, Freiburg, Germany), was equilibrated with a buffer solution containing 100 mM NaCl, 0.5 mM EDTA, 20 mM Tris-HCl, pH 7.5. The solution containing the freshly prepared NDs were applied to the column and purified through Äkta (GE Healthcare Life Sciences) fast protein liquid chromatography (FPLC), at a flow rate of 0.4 ml/min and dual absorbance at wavelength of 280 nm and 215 nm. The peaks containing NDs were collected and concentrated using a 10 kDA molecular weight cutoff.

Nanodiscs prepared using MSPD1ΔH5 have an overall diameter of approximately 8 nm, with an inner diameter of about 6 nm and a total surface area of roughly 2826 Å². These medium-sized nanodiscs remain stable for at least 30 days when stored at 4 °C.

### 4.4. Dynamic Light Scattering

Dynamic Light Scattering (DLS) measurements were carried out using Malvern Zetasizer Nano S (Malvern Instruments, UK) equipped with a 4 mW He−Ne gas laser (beam wavelength = 632.8 nm) and 173° back scattering measurement facility. Measurements were taken using a low-volume disposable cuvette. The Z-average diameter was calculated from the correlation function using Malvern technology software.

### 4.5. Transmission Electron Microscopy

10 µL of each sample was spotted on a hydrated carbon-coated copper grid (Electron Microscopy Sciences, USA) and left for 2 min, then blotted to remove excess buffer. Subsequently, the samples were washed with milli-Q water and stained with a freshly prepared, filtered 1 % (w/v) uranyl acetate solution (Electron Microscopy Sciences, USA) for 5 min. Finally, samples were air-dried overnight in a dust-free space. A transmission electron microscope (FEI Tecnai TF20) operating at 200 kV with a point resolution of 0.24 nm and a line resolution of 0.102 nm was used for imaging. Digital image recording was used, and EDS mode was used for analysis. The width of the fibrils of Aβ40 mutants (G33L and G37L) in the absence and presence of ND were analysed using the Image J software.

### 4.6. Circular Dichroism spectroscopy

A JASCO J-1500 CD spectrometer (Japan) was used to measure the CD utilizing a 0.1 cm path length cell with a 2 nm slit width. 20 mM phosphate buffer (pH 7.3) containing 50 mM NaF was used to dissolve peptides at a concentration of about 80 µM. At 25 °C, far-UV CD spectra were recorded between 260 and 190 nm, at a scan rate of 100 nm/min. Three measurements, or three accumulations, were obtained for each spectrum, and the average was taken. The raw data were processed using buffer subtraction and smoothing.

### 4.7. NMR experiments

NMR experiments were performed on a Bruker Avance III 500 MHz NMR spectrometer equipped with a 5 mm SMART probe and a Bruker Avance III 700 MHz spectrometer equipped with a RT probe. All NMR experiments were performed either at 10 °C or 25 °C. A two-dimensional ^1^H-^15^N heteronuclear quantum coherence (HSQC) NMR spectrum of monomeric Aβ40 was recorded. The pulse program used was hsqcetf3gpsi (Table S2). The resonance assignments of the monomer obtained from 2D ^1^H-^15^N HSQC spectra was done following our previous work (38). The fibril was then added to the monomeric solution in NMR tube, to enhance the aggregation kinetics. To monitor the effect on Aβ40 aggregation, in presence of NDs (L/P=20) at atomic resolution, real-time 2D NMR experiments were performed by consecutively acquiring ^1^H-^15^N HSQC spectra in the absence and presence of NDs. Each spectrum was obtained from 64 t_1_ experiments, 16 scans, 8 dummy scans, and a 1 s recycle delay. The points collected in proton dimensions is 1024.The spectral widths were set at 16 and 40 ppm, and the offsets were 4.7 and 117 ppm for the proton and nitrogen dimensions, respectively.

For AV20, 2D total correlation spectroscopy (TOCSY) NMR experiments were performed at 25 °C on the Bruker Avance 700 MHz spectrometer. Real-time 2D HSQC NMR experiments were performed for AV20 with ^15^N-labeled Gly residues, on the Bruker Avance III 500 MHz NMR spectrometer at 10 °C.

The two-dimensional (2D) ^1^H-^15^N HSQC experiments were performed at 25 °C with an array of delays of 0, 14.4, 28.8, 43.2, 53.2, 72, 100.8, 129.6, and 158.4 ms with a repetition at 72.0 ms. For all the experiments, a total of 1024 (t_2_) and 64 (t_1_) data points were collected, and the number of scans was 16. hsqct2etf3gpsi pulse program was used with Echo/Antiecho-TPPI gradient selection. The Origin software (Origin 2018) was used to fit the data using mono-exponential equation to obtain R_2_ values.

^1^H-^15^N CEST experiments were performed based on the published pulse sequence (59). All CEST experiments were performed at 25 °C. A series of 22 two-dimensional datasets were acquired at echo-Antiecho mode, corresponding to ^15^N offsets between 104 ppm and 130.3 ppm obtained at 50 Hz (0.7 ppm) interval. The TD values were 2096 (t_2_) and 256 (t_1_) and number of scans was 16. The data was processed and analysed using Bruker Topspin 3.7.0. and Bruker Dynamic Centre version 2.8.6.

Triple resonance NMR experiments i.e., HNCA and CBCACONH were performed with the sample of Aβ40 co-incubated with nanodisc after 48 h of incubation on a Bruker Avance III 700 MHz spectrometer equipped with a 5 mm cryogenic probe at 25 °C. For HNCA and CBCACONH, dummy scans were 32, spectral widths were 12, 36 and 40 ppm for the proton, nitrogen and carbon dimensions, respectively, Offsets were 4.69, 117 and 54 ppm for the proton, nitrogen and carbon dimensions, respectively, TD values were 2048, 40 and 128 for the proton, nitrogen and carbon dimensions, respectively. For HNCA, the spectral widths were 16, 36 and 40 ppm, for the proton, nitrogen and carbon dimensions, respectively. For CBCACONH, the spectral widths were 16, 36 and 80 ppm, for the proton, nitrogen and carbon dimensions, respectively. The obtained data were processed using nmrPipe and nmrDraw data processing and analysis suites (http://spin.niddk.nih.gov/NMRPipe/) on a Linux based computer system. Further analysis and peak assignment were done in SPARKY 3.113 (https://www.cgl.ucsf.edu/home/sparky). The pulse programs for all the NMR experiments are given in Supporting information Table S2.

### 4.8. Coarse-grained (CG) molecular dynamics (MD) simulations

For preparing the initial system comprising ten molecules of Aβ40 and ND, the solution NMR structure of Aβ40 (PDB ID: 2LFM) was used (60). Ten Aβ40 molecules were placed randomly more than 5 Å away from each other. The ND comprised of two MSP molecules (marked as MSP-1 and MSP-2 with same sequence) and DLPC and DLPG lipids at 3:1 ratio was prepared using CHARMM-GUI (61). DLPC and DLPG were chosen for CG MD simulation, because the 12-carbon and 14-carbon lipid tails containing DLPC/DLPG and DMPC/DMPG respectively, are represented by the identical CG models (DLPC/DLPG) (62). Besides that, the primary aim of the study is the interaction of Aβ40 with the lipid head group; the lipid tails have no direct involvement in the interaction with Aβ40. The Aβ40 molecules are present more than ∼ 8 Å away from the lipid layer of the ND. The system was neutralized with counter ions, besides maintaining a concentration of 0.15 M NaCl. The simulation was performed with Martini force field (63), the corresponding coarse-grained water model was used for the solvation of the system. After energy minimization and equilibration process at 310 K, the final production run was performed for 2 μs in triplicate. GROMACS 2023 software (64) was used for performing all the simulation processes. The final trajectories were analyzed using GROMACS and visual molecular dynamics (VMD) (65) and figures were prepared using PyMol software.

For preparing the initial structure of Aβ40G33L and Aβ40G37L mutants, the Gly residues were mutated to Leu residues using Coot (48) in the wild-type Aβ40 structure. The CG simulations of Aβ40G33L and Aβ40G37L mutants were performed in a similar fashion to the wild type consisting of ten monomers in the same orientation as the wild-type. The overall simulation process was performed in the same way as it was performed for the wild-type. In the case of AV20, the twenty residues from the N-terminal were deleted from the wild-type structure. The CG simulation was performed using the same process as mentioned for the wild-type, consisting of ten monomers in the same orientation. The final production runs of the simulations for AV20, Aβ40G33L and Aβ40G37L mutants were also performed for 2 μs in triplicate.

### 4.9. All-atom MD simulations

The Aβ40 structure (PDB ID: 2LFM) used for the CG simulation is also used for the all-atom simulation process. The model used for the all-atom simulation process also consists of ten monomers in a similar orientation to the CG model. The protonation state of the Aβ40 monomers was defined based on pH 7.4. The ND contains two MSP molecules along with DMPC and DMPG lipids at a 3:1 ratio, prepared using CHARMM-GUI. The whole system was solvated using the TIP3P water model and NaCl was added at a concentration of 0.15 M. The simulation was performed with AMBER14SB force field (66) using GROMACS 2022 software. Following energy minimization of the system, the NVT equilibration was performed using V-rescale thermostat at 310 K, after which NPT equilibration was done using C-rescale barostat. The final production run was performed for 150 ns with the same thermostat and barostat. Electrostatic and Van Der Waals interactions between the atom pairs were calculated using particle mesh Edward method and Verlet scheme with cut-off of 1 nm for both the interactions during the simulation process.

### 4.10. Statistical Analysis

All experiments were performed in three replicates. All statistical analysis was performed in Graph Pad Prism 8. Results are expressed as mean± SEM. Statistical significance was determined using the two-tailed, unpaired Student’s t-test in column analysis or one-way ANOVA in group analysis. p ≤ 0.05 (95% confidence) was considered significant in all studies.

## Data availability

The raw data are provided as a separate file in the Supporting Information.

## Supporting Information

This article contains supporting information. The figures supporting the main manuscript are provided in the Supporting Information.

## Author Contributions

AB and DS conceived the project, and designed the research; AB arranged the funding for this work; DS performed all biophysical assays, solution-state NMR experiments; DR expressed and purified the nanodiscs, did 3D NMR assignments, relaxation experiments and CEST with JK; CS helped in expressing and purification of nanodiscs to DR; SC performed CG and all atom MD simulation under the supervision of PB; NM, RB and DulalS did all the TEM analysis; NS procured the peptides and edited the manuscript; SM performed size exclusion chromatography (SEC) and ThT experiments with DG; DS, KB, JK, DR and AB analyzed solution-state NMR data; DS, DR, SC, PB, KB, DulalS and AB wrote the manuscript; all authors reviewed the manuscript. ‡DS, ‡DR, ‡KB contributed equally.

## Funding and additional information

This work was partly supported by Department of Biotechnology (BT/PR29978/MED/30/2037/2018 and (BT/54681/BMS2/156/82/2024 to AB), Government of India, partly by Council of Scientific and Industrial Research (02(0292)/17/EMR-II to AB), Government of India to AB. DS thanks DST-INSPIRE for providing SRF and DR acknowledges CSIR-UGC for SRF. KB acknowledges Velux Stiftung, Switzerland for research associate position. The authors acknowledge central instrument facilities (CIF) of Bose Institute.

## Funding and additional information

The authors declare that they have no conflicts of interest with the contents of this article.

## ABBREVIATIONS

Aβ: Amyloid Beta;
AD: Alzheimer’s Disease;
MSP: Membrane Scaffold Protein;
ND: Nanodisc;
SUV: Small Unilamellar Vesicles;
CD: Circular Dichroism;
NMR: Nuclear Magnetic Resonance;
ThT: Thioflavin T;
HSQC: Heteronuclear Single Quantum Coherence;
MD: Molecular Dynamics;
CEST: Chemical Exchange Saturation Transfer.

