## Supporting Information for "Nanodisc-Enabled Dissection of GxxxG- Mediated Amyloid Beta Oligomerization Mechanisms"

**Table S1:** Details of NDs used in this study.

| Lipid Mixture | MSP Variant | MSP/Lipid Ratio | Temperature applied during NDs assembly (°C) |
|---|---|---|---|
| 75 % DMPC<br>25 % DMPG | MSPD1ΔH5 | 1: 50 | 30 |

**Table S2:** The pulse programs (Bruker) used for the NMR experiments in the study.

| **NMR experiment** | **Pulse programme** |
|---|---|
| $T_2$ relaxation | *hsqct2etf3gpsi* |
| $T_1$ relaxation | *hsqct1etf3gpsi.2* |
| Chemical Exchange Saturation Transfer (CEST) | *hsqc_cest_etf3gpsitc3d* |
| $^1H$-$^{15}N$ Heteronuclear Quantum Coherence (HSQC) | *hsqcetf3gpsi* |
| 3D HNCA | *hncagp3d* |
| 3D HNCACB | *hncacbgp3d* |
| 3D CBCACONH | *cbcaconhgp3d* |
| 2D TOCSY | *mlevesgpph* |

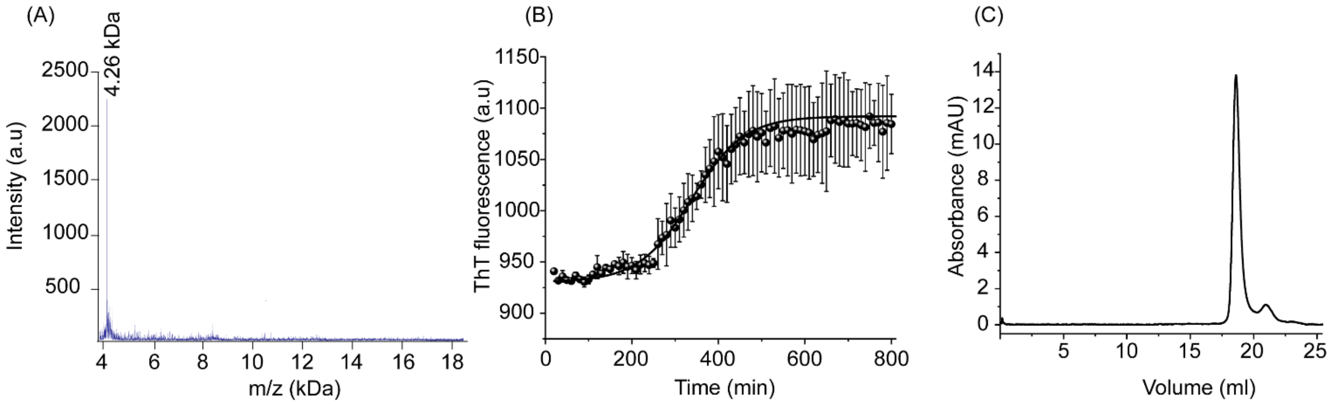


**Figure S1:** (A) MALDI analysis of freshly prepared Aβ40 peptide showing a single peak at 4.26 kDa. (B) ThT fluorescence assay of the 20 μM Aβ40 peptide under shaking conditions (250 rpm). Each point represents the mean of three replicates, with error bars indicating SD. (C) SEC profile of the freshly prepared Aβ40 peptide showing a single peak at ~20 ml. The experiments were repeated thrice.

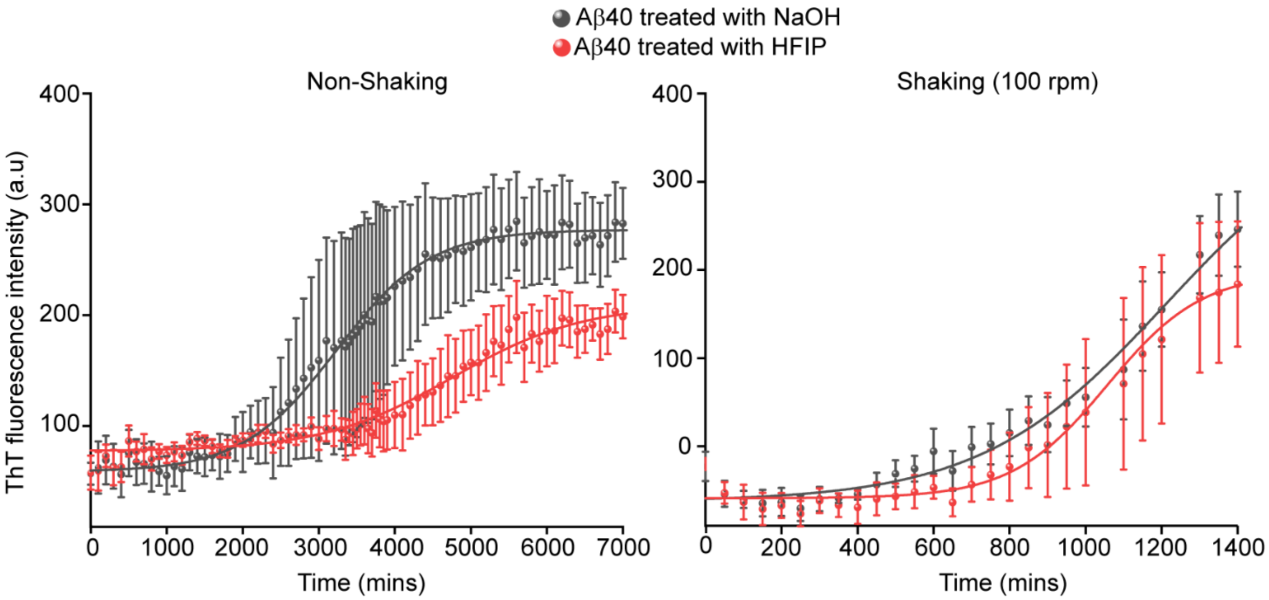


**Figure S2:** The effect of treatment of NaOH and HFIP on the aggregation of Aβ40 under non shaking (left panel) as well as shaking (100 rpm) conditions (right panel).

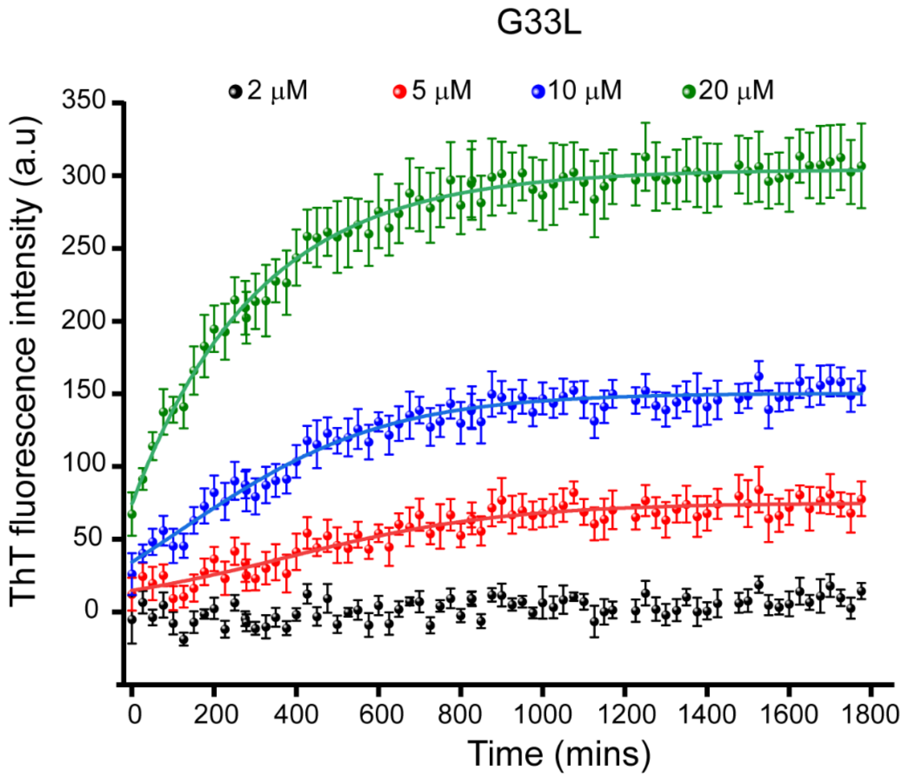


**Figure S3:** Concentration-dependent aggregation kinetics of the Aβ40G33L mutant monitored using ThT fluorescence. At 2 μM, there was no increase in fluorescence, indicating no aggregation. At 5 μM, the increase in fluorescence intensity indicates aggregation of Aβ40G33L. No lag phase was observed across the concentration range of 5–20 μM, indicating that the mutants do not follow a sigmoidal curve like wild-type Aβ40 peptide.

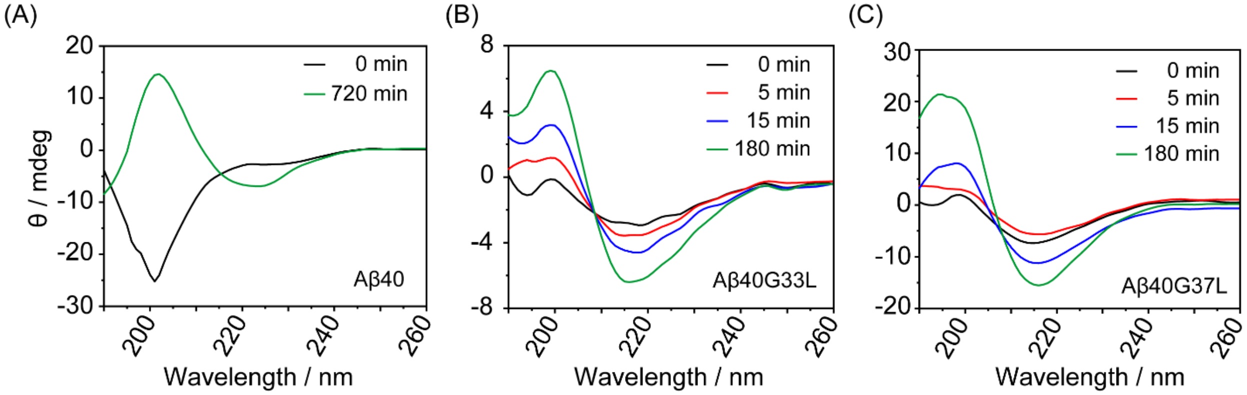


**Figure S4.** Far-UV CD spectra of (A) Aβ40, (B) Aβ40G33L, and (C) Aβ40G37L at different time-point showing the time-dependent structural transition from monomer to fibril. All samples contain ~80 μM peptide in 20 mM sodium phosphate (pH 7.4) and 50 mM sodium fluoride, incubated at 37 °C, and 250 rpm shaking conditions. All experiments were repeated three times.

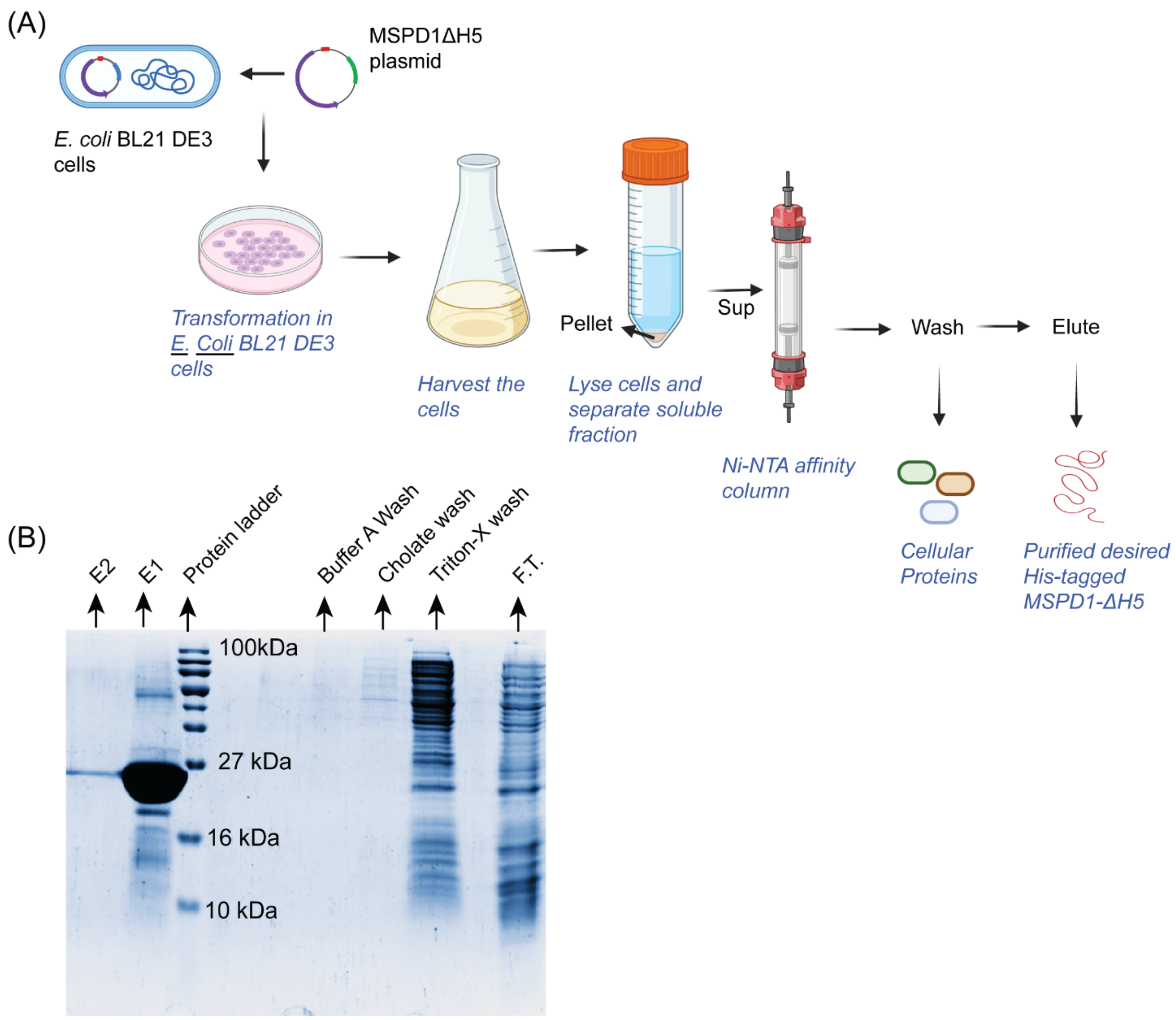


**Figure S5.** (A) Illustration of the steps involved in the overexpression of MSP in *E.coli* BL21 cells and purification using Ni- NTA affinity column created in BioRender.com. (B) Image of the SDS PAGE of samples obtained at various steps of MSP purification. The lanes contain Flow through (F.T.), Triton-X wash, Cholate wash, Buffer A wash Protein ladder, elute 1 (E1) and elute 2 (E2), from right to left.

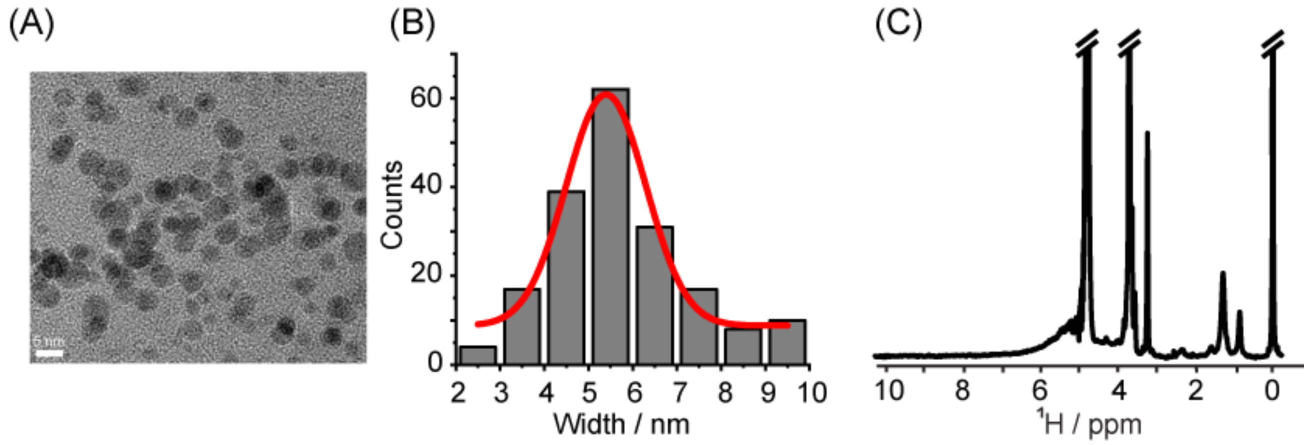


**Figure S6.** Characterization of NDs. (A) HR-TEM image, (B) Count vs Width plot derived from TEM images and (C) 1D $^{1}$H NMR of free NDs. The experiment was performed using Bruker 500 MHz NMR spectrometer and at 25 °C.

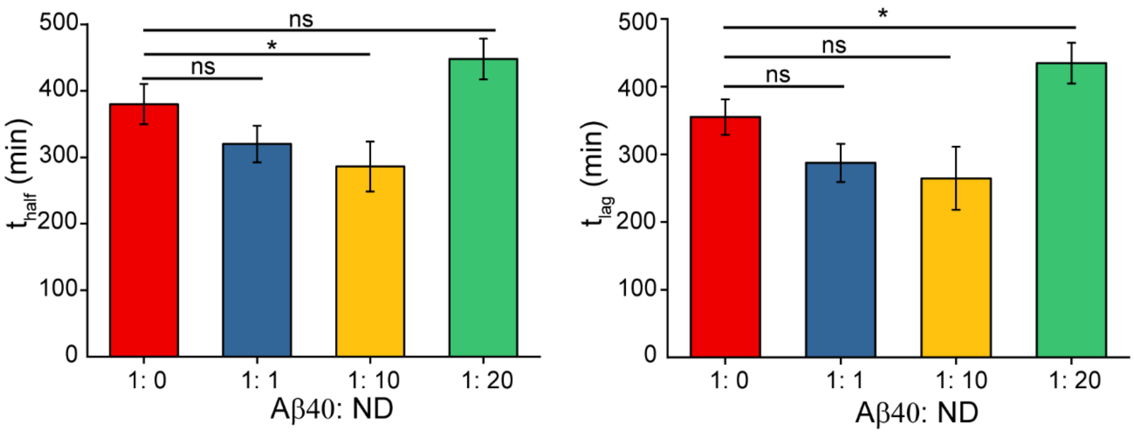


**Figure S7.** Aggregation half-life ($t_{half}$) and lag time ($t_{lag}$) calculated from the curve fit using a sigmoidal growth model. Each column represents the mean of three replicates, with error bars indicating SEM. Significant changes are marked with * ($p \leq 0.05$) according to one- way ANOVA test, while "ns" indicates non-significance.

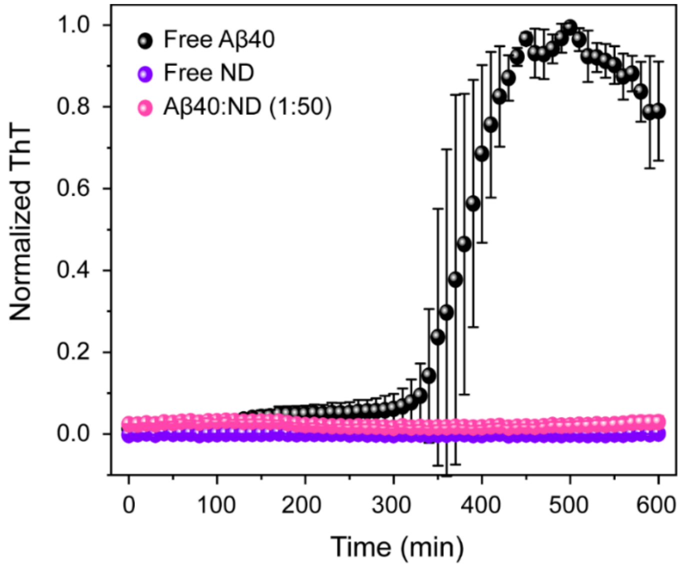


**Figure S8.** Aggregation kinetics of Aβ40 (20 μM) in presence of nanodiscs (ND) at 1:50 peptide-lipid ratios (L/P=50) showing a drastic increase in the lag time in the presence of such high concentration of lipid. No increase in the ThT intensity was observed for nanodiscs (200 μM) alone. Fluorescence measurements were taken in a 96 well plate under shaking (300 rpm, 30s before each cycle 120s) at 37 °C. All experiments were repeated three times, and the data was averaged (±SD).

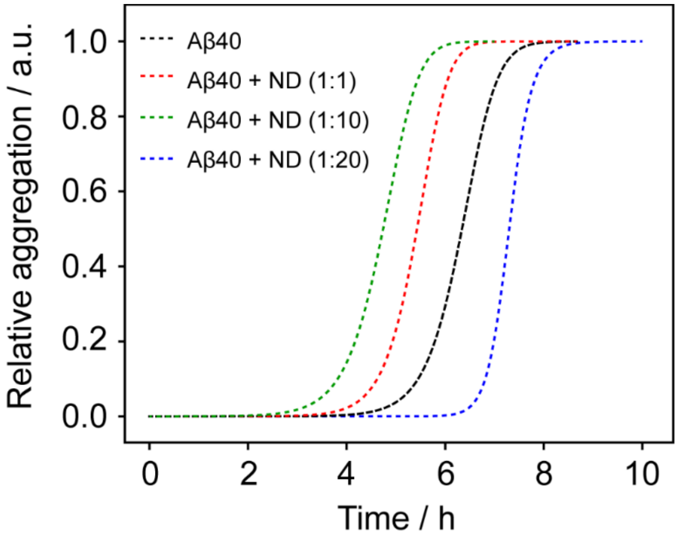


**Figure S9.** Fitting of the experimental aggregation-kinetics data of Aβ40 in the absence and presence of NDs at different peptide: lipid ratios using the multi-step secondary nucleation model in AmyloFit.(2)

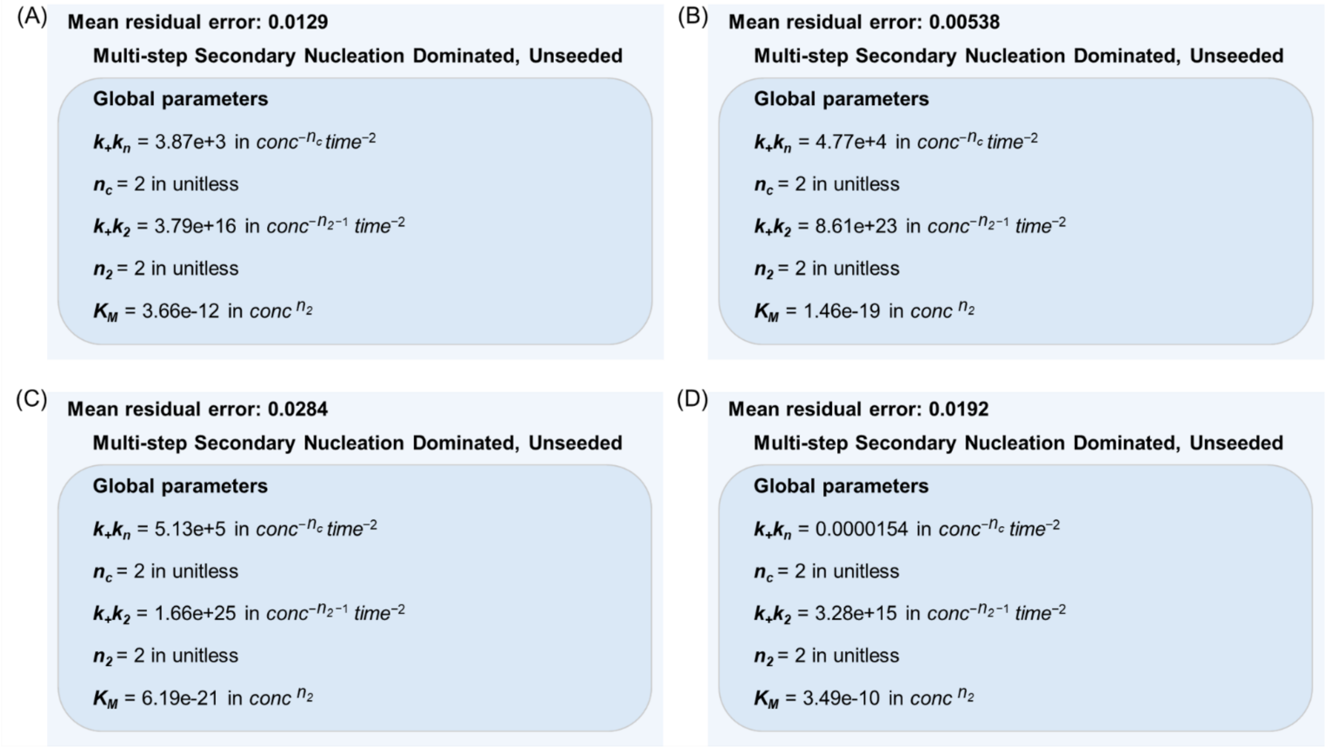


**Figure S10.** Parameters obtained from the global fitting of (A) free Aβ40, (B) Aβ40 + NDs (1:1), (C) Aβ40 + NDs (1:10), and (D) Aβ40 + NDs (1:20) kinetic ThT data using multi-step secondary nucleation model. According to this model, the first step—where single peptide units attach to the fibril surface—depends on how many monomers are present. After that, steps that don't rely on monomer concentration take place. These may involve the monomers detaching from the fibril, forming a nucleus, and changing their shape. The rate constants for elongation ($k^+$), primary nucleation ($k_n$), secondary nucleation ($k_2$), and the Michaelis constant ($K_m$) were all calculated by fitting the model to the data.

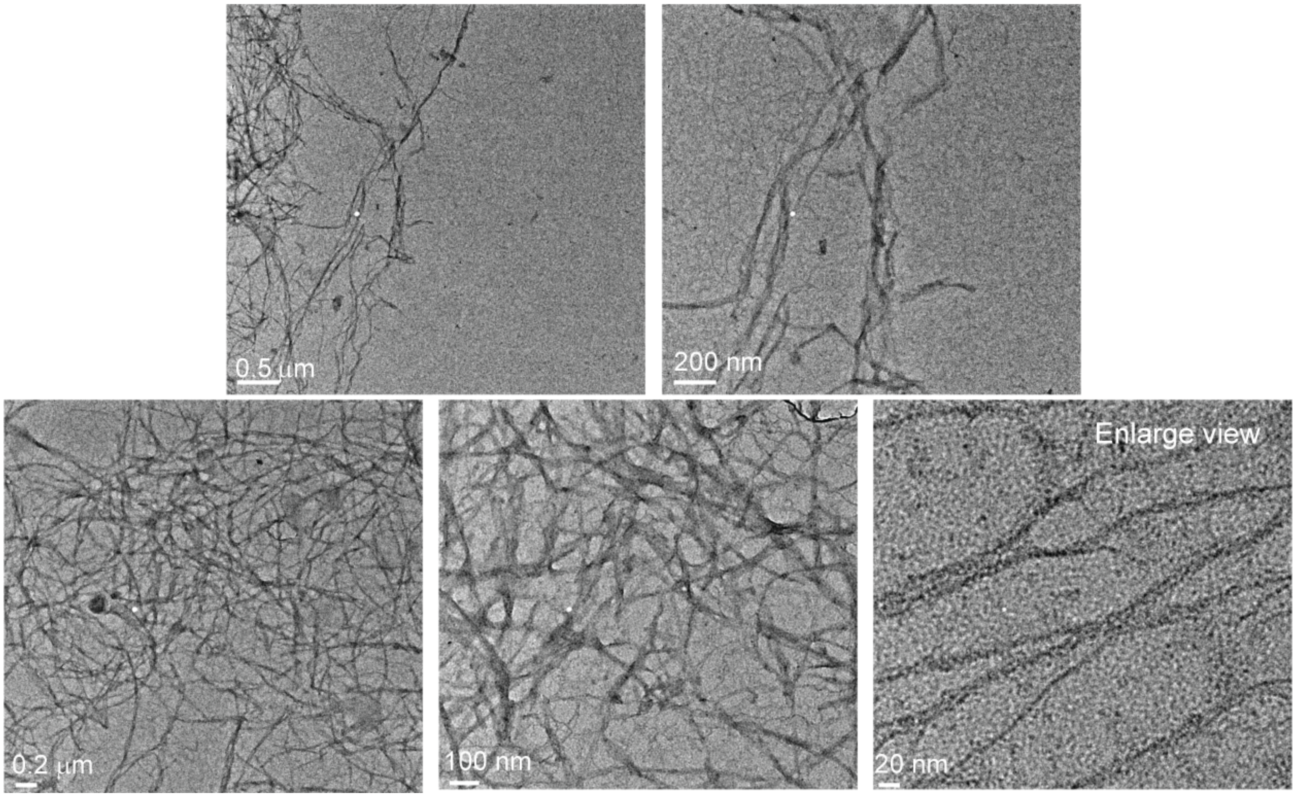


**Figure S11.** HR-TEM images of Aβ40 fibers at different magnifications showing dense network of amyloid fibers of different width.

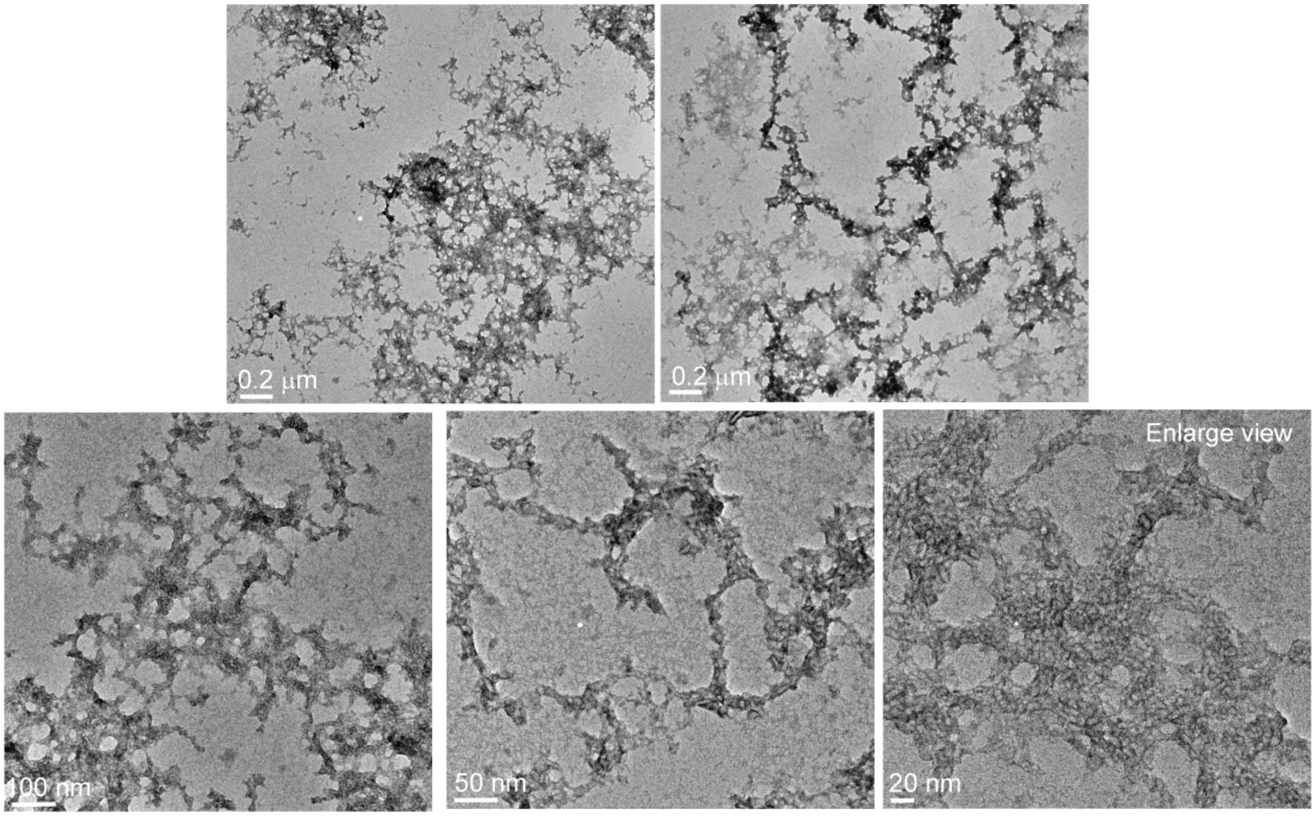


**Figure S12.** HR-TEM images of Aβ40 in presence of nanodiscs (L/P=20) different magnifications showing disordered aggregates. No dense network of amyloid fibers was detected.

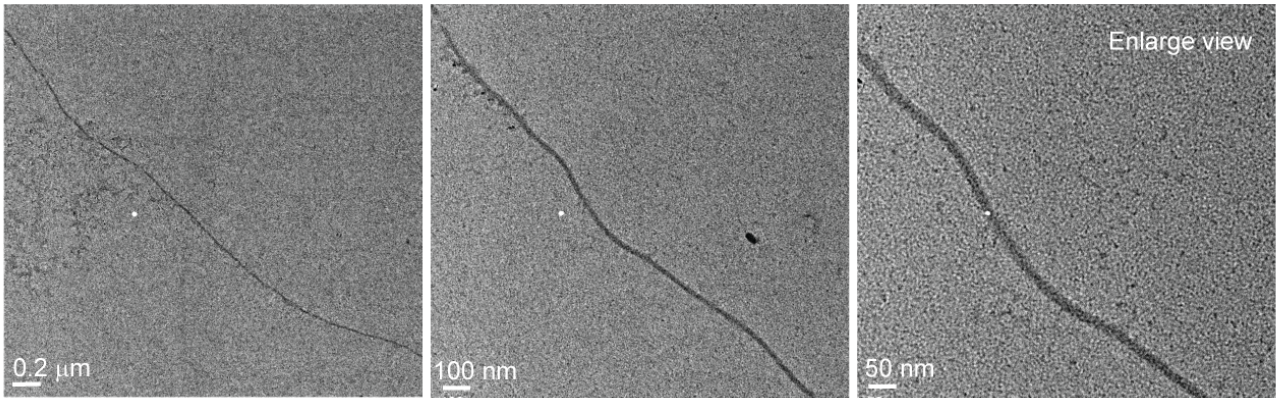


**Figure S13.** HR-TEM images of Aβ40G33L fibers at different magnifications showing long isolated fibers.

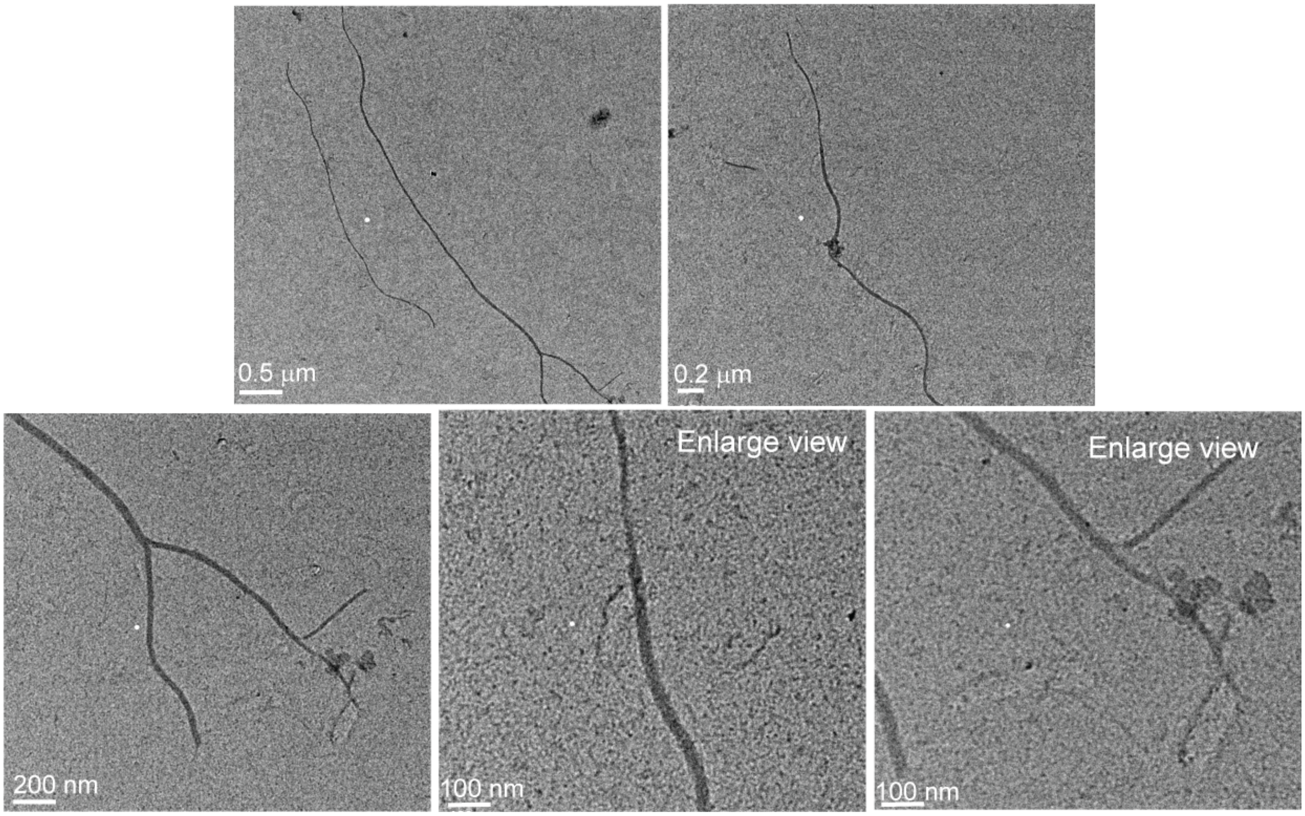


**Figure S14.** HR-TEM images of Aβ40G33L in presence of nanodiscs (L/P=20) at different magnifications showing long isolated fibers.

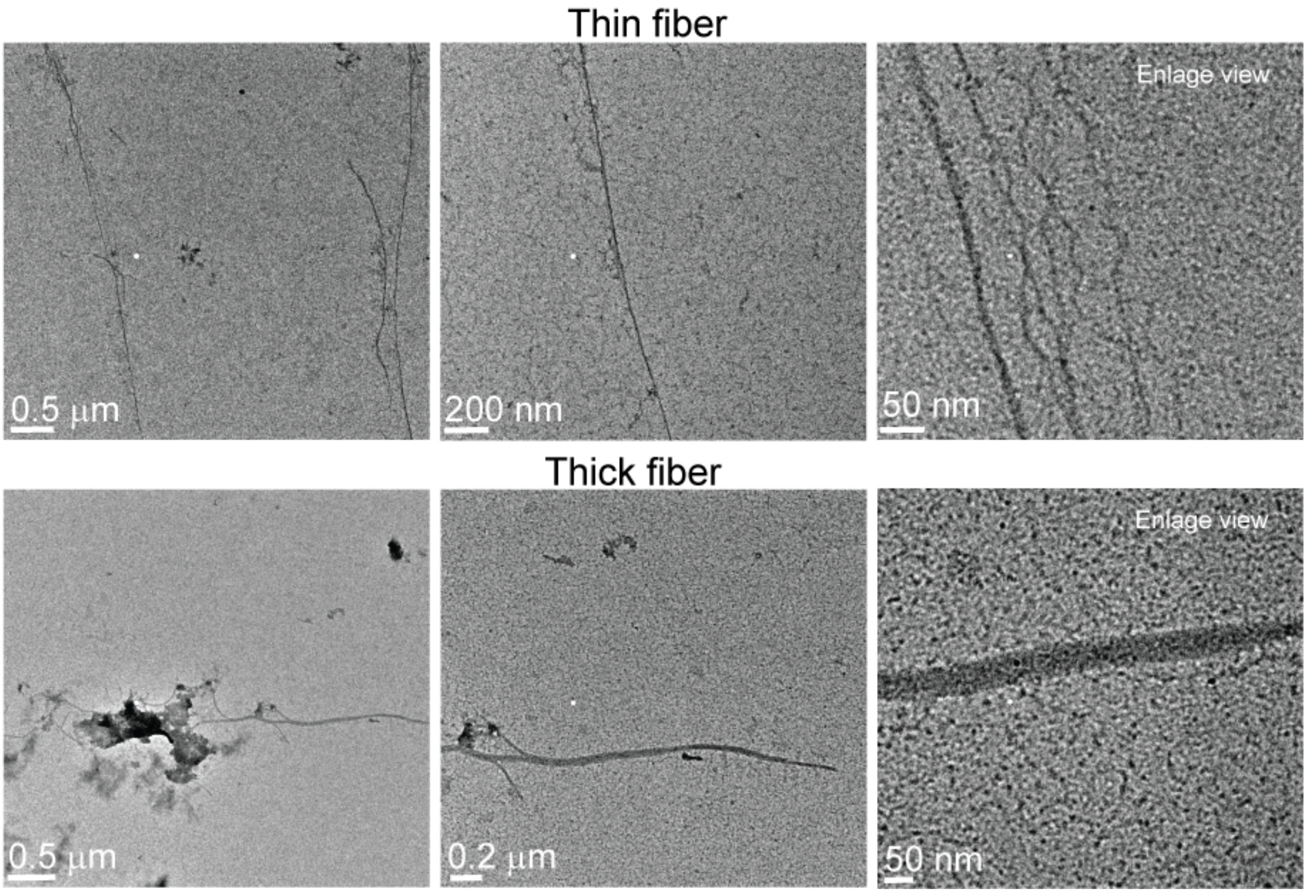


**Figure S15.** HR-TEM images of Aβ40G37L fibers at different magnifications showing long isolated fibers of different width.

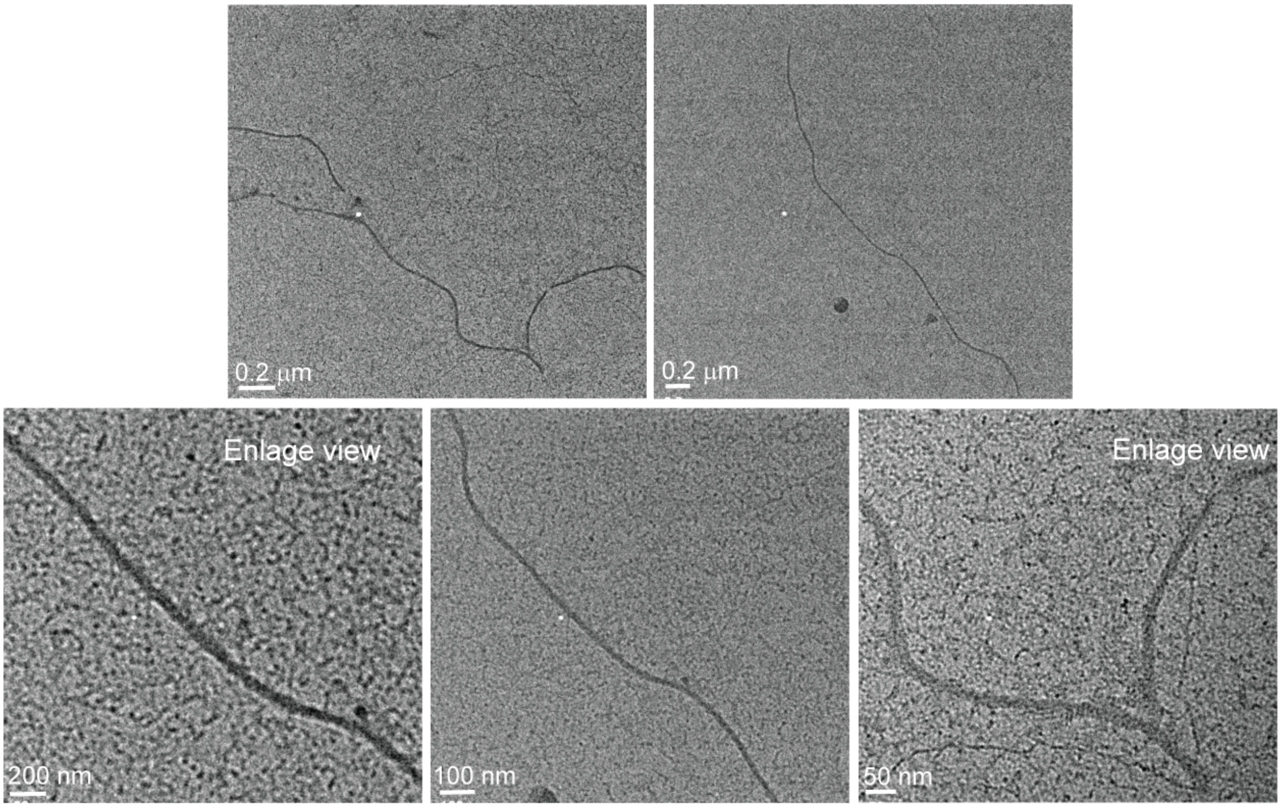


**Figure S16.** HR-TEM images of Aβ40G37L in presence of nanodiscs (L/P=20) at different magnifications showing long isolated fibers.

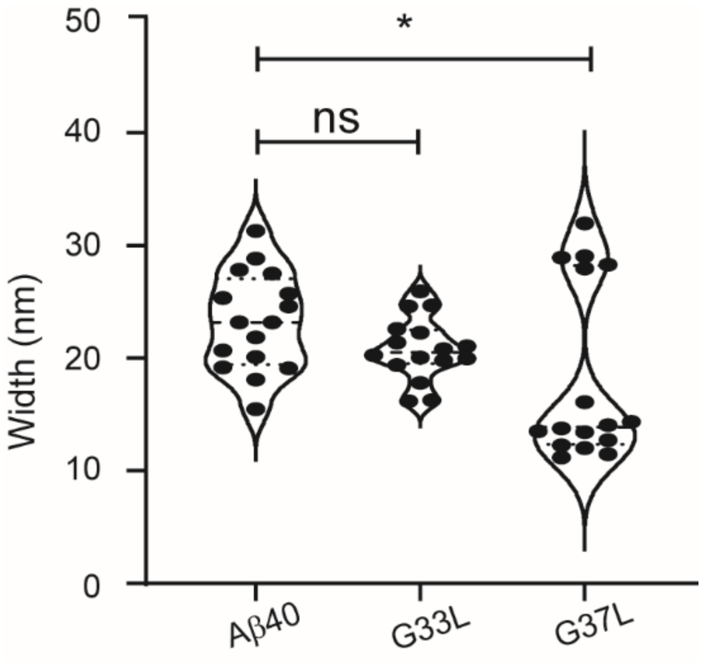


**Figure S17.** Analysis of the width of the fibrils of Aβ40 peptides and the mutants (G33L and G37L) using Image J software. Aβ40 peptides as well as the mutants formed fibrils with diverse width size. Significant changes are marked with *($p \leq 0.05$) according to the two-tailed, unpaired Student's t-test, while "ns" indicates non-significance.

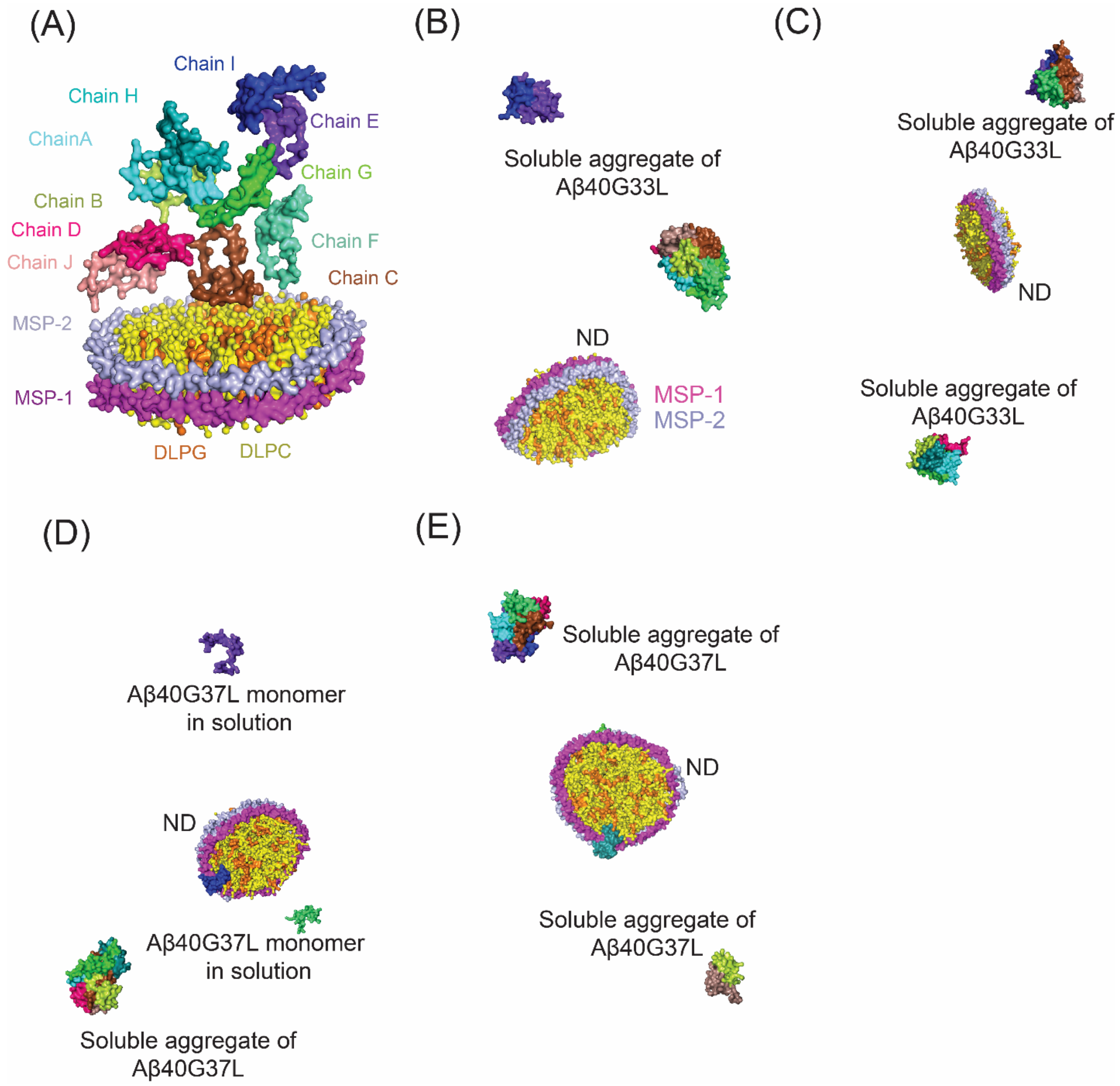


**Figure S18.** (A) Initial structure of NDs containing MSP proteins and DLPC and DLPG lipid models and ten monomers of Aβ40 used for coarse-grained (CG) MD simulation. The DLPC and DLPG molecules are shown in yellow and orange cartoon representations, respectively. The two MSP molecules are shown in magenta and light blue surface representations. Aβ40 chains are shown in multiple colors in surface representation. Formation of soluble aggregate (B), (C) Aβ40G33L, (D), (E) Aβ40G37L without formation of aggregate on NDs as observed in multiple simulations run for 2μs.

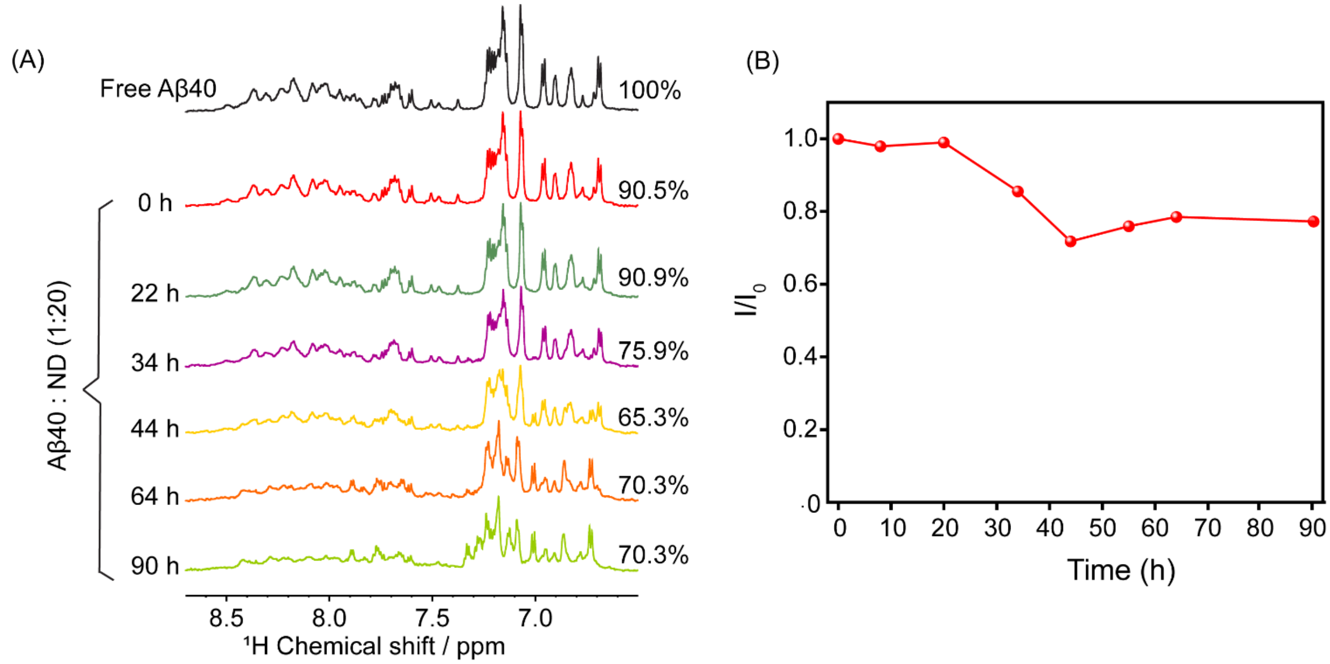


**Figure S19.** (A) Selected one-dimensional $^{1}$H NMR spectra of Aβ40 in the absence and presence of NDs (L/P=20) at different time points at room temperature. Upon addition of NDs to Aβ40, the amide proton signal intensity decreased by ~10% at 0 h. The decrease in amide proton signal intensity reached 35 % at 44 h, following a significant chemical-shift perturbation. (B) Plot showing the decay in amide proton signal intensity ($I/I_0$) of Aβ40 in the presence of NDs (L/P=20) as a function of time.

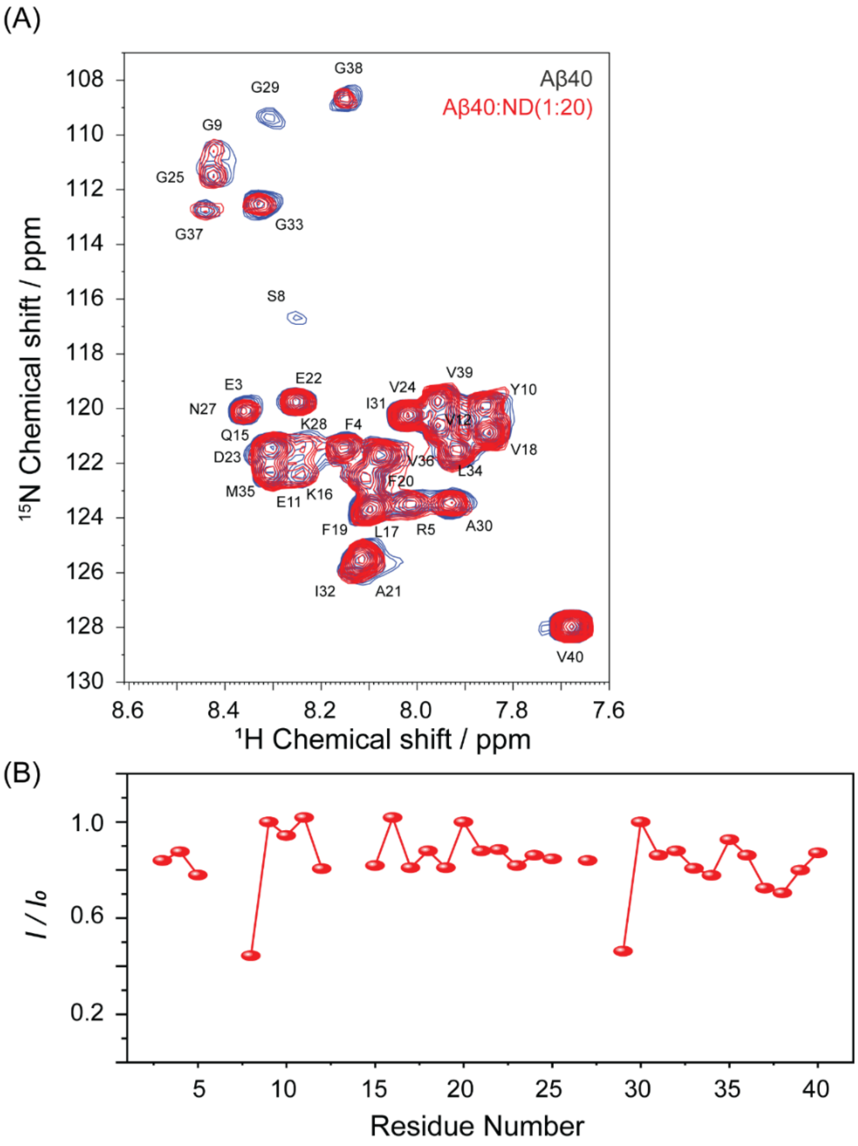


**Figure S20.** (A) $^{15}$N HSQC overlay spectra of Aβ40 in absence (blue) and presence of nanodiscs (L/P=20) at room temperature, suggesting slight intensity drop without any notable chemical shift perturbations. While it is correct that a few resonances of the free Aβ40 peptide (e.g., S8 and G29) exhibit relatively lower signal-to-noise (S/N) ratios, the intensity ratio analysis is based on the overall residue-wise trend rather than isolated low-intensity peaks, and this trend is consistently observed across multiple residues with good signal-to-noise ratios. (B) Changes in the peak intensities along the peptide backbone measured from 2D $^{1}$H-$^{15}$N HSQC spectra upon titration of a freshly prepared Aβ40 (~80 μM) sample with nanodiscs (L/P=20) at 25 °C. Points represent the signal intensity of the peak maxima (*I*) relative to the signal intensity in the absence of nanodiscs ($I_0$). Residues with lower signal-to-noise may carry higher uncertainty but do not impact the overall conclusions.

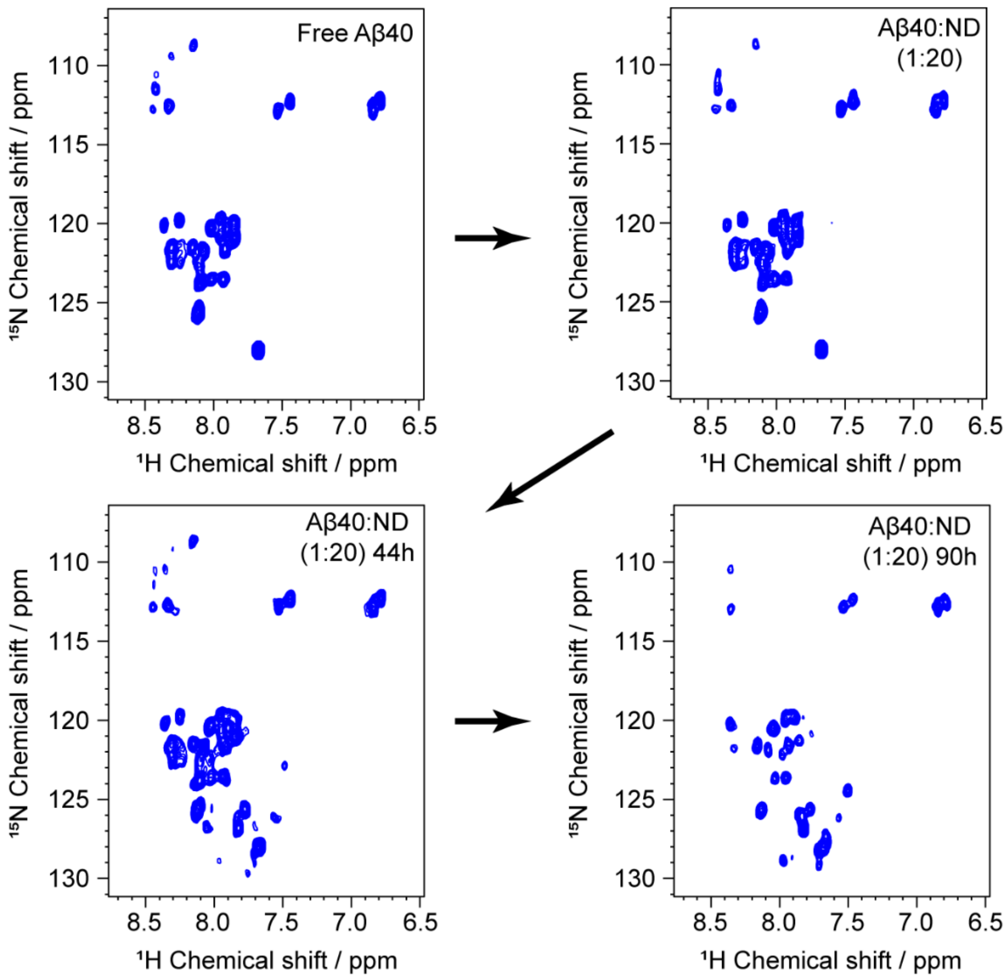


**Figure S21.** Representative real-time HSQC spectra of Aβ40 (80 μM) aggregation in presence of nanodiscs (L/P=20) at room temperature. HSQC spectra were acquired sequentially at 700 MHz over a period of 100 h at 25 ºC in 50 mM NaCl, 20 mM phosphate buffer, pH 7.4.

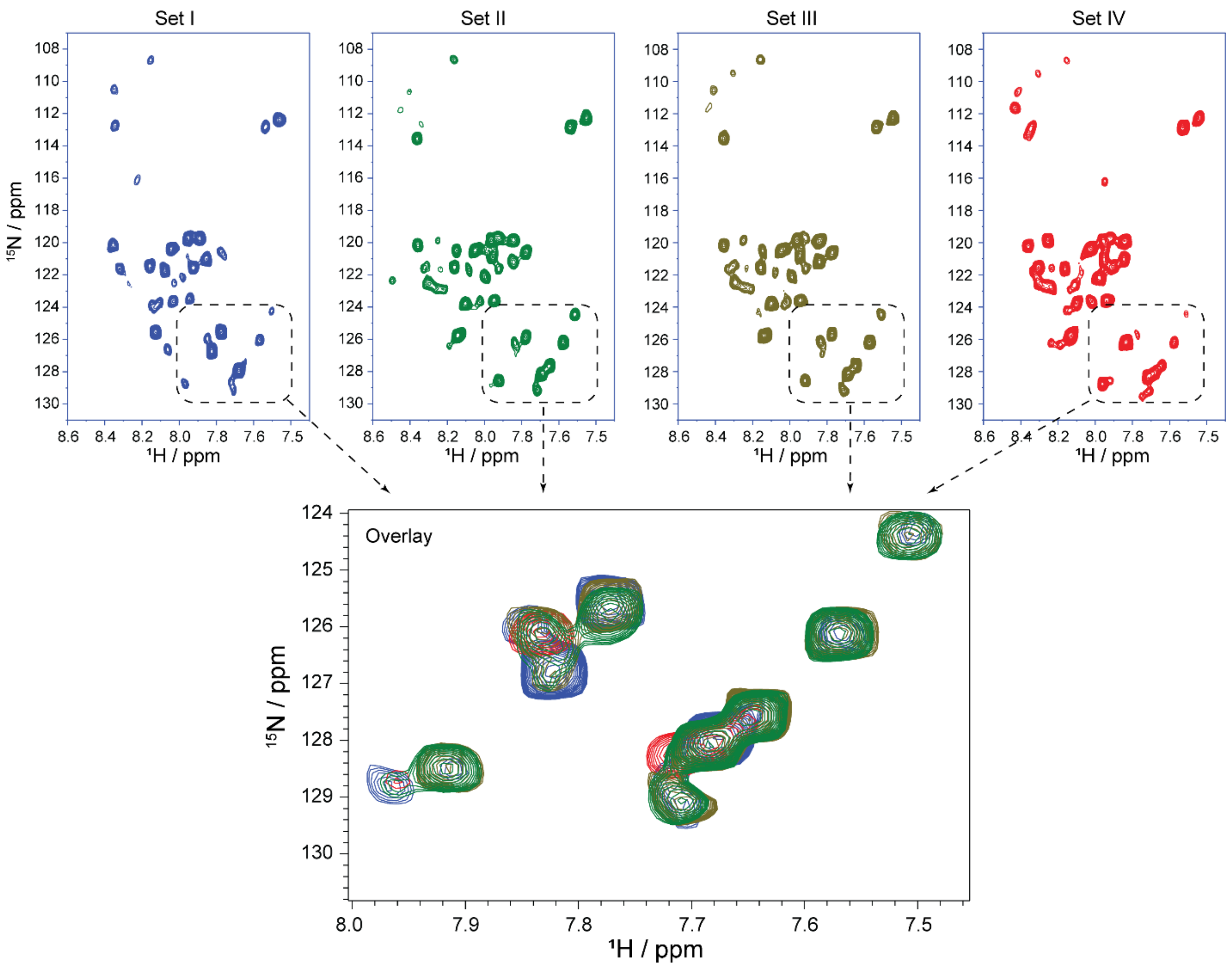


**Figure S22.** $^{15}$N HSQC spectra of Aβ40 (~80 μM) aggregates in presence of nanodiscs (L/P=20) at room temperature, performed for multiple batches, showing generation of similar new peaks in the vicinity of V40. The experiments were performed using Bruker 700 MHz NMR spectrometer.

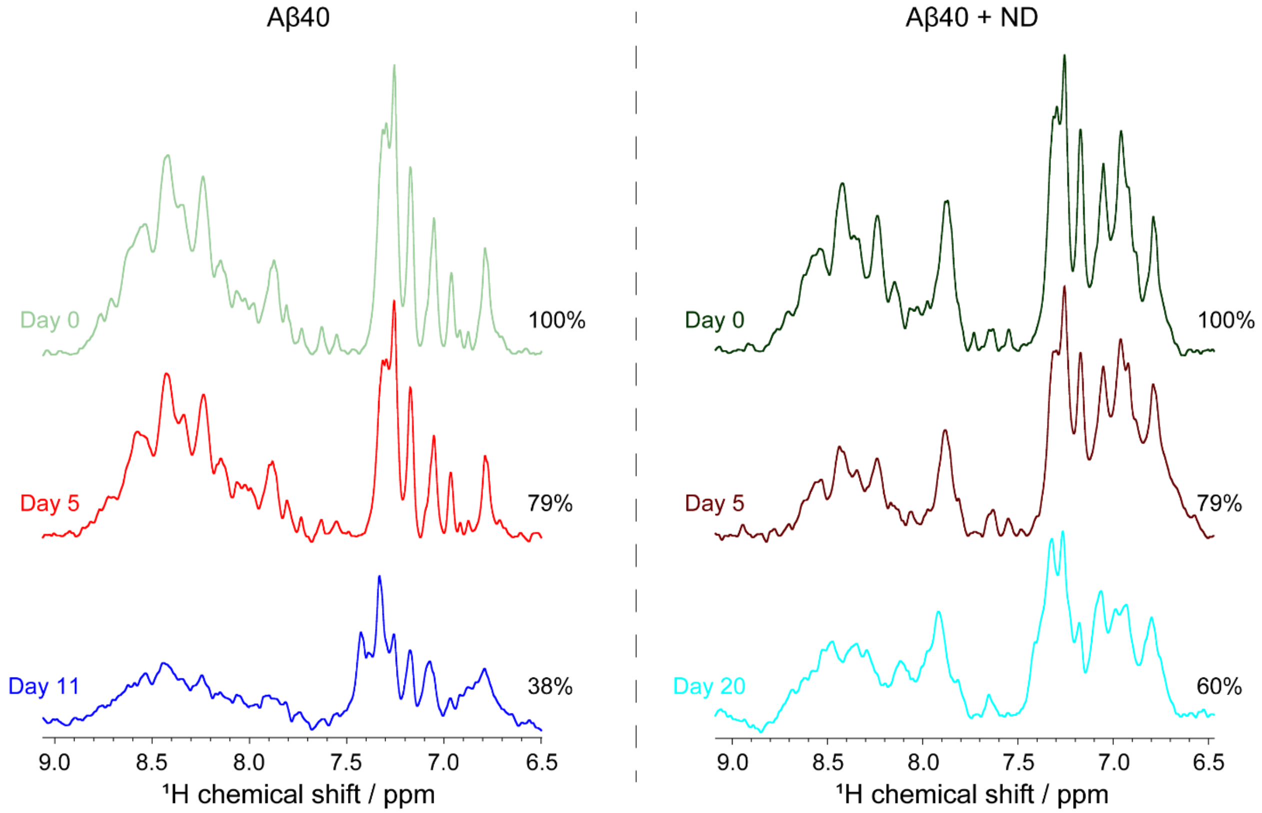


**Figure S23.** 1D $^{1}H$ NMR spectra of Aβ40 in absence and presence of nanodiscs at different time intervals. Samples were incubated at 25 °C under non-shaking conditions and spectra were taken at 10 °C. using Bruker Avance 700 MHz NMR spectrometer. Initially both Aβ40 and Aβ40 in presence of high lipid concentration showed almost 20 % signal decay at day 5. After this time point, Aβ40 underwent drastic decrease in the signal intensity indicating fibrillation, whereas Aβ40 in presence of nanodiscs showed a slow decrease in the signal intensity. Even at 20 days of incubation, sufficient NMR signal was observed suggesting prolongation of oligomeric states.

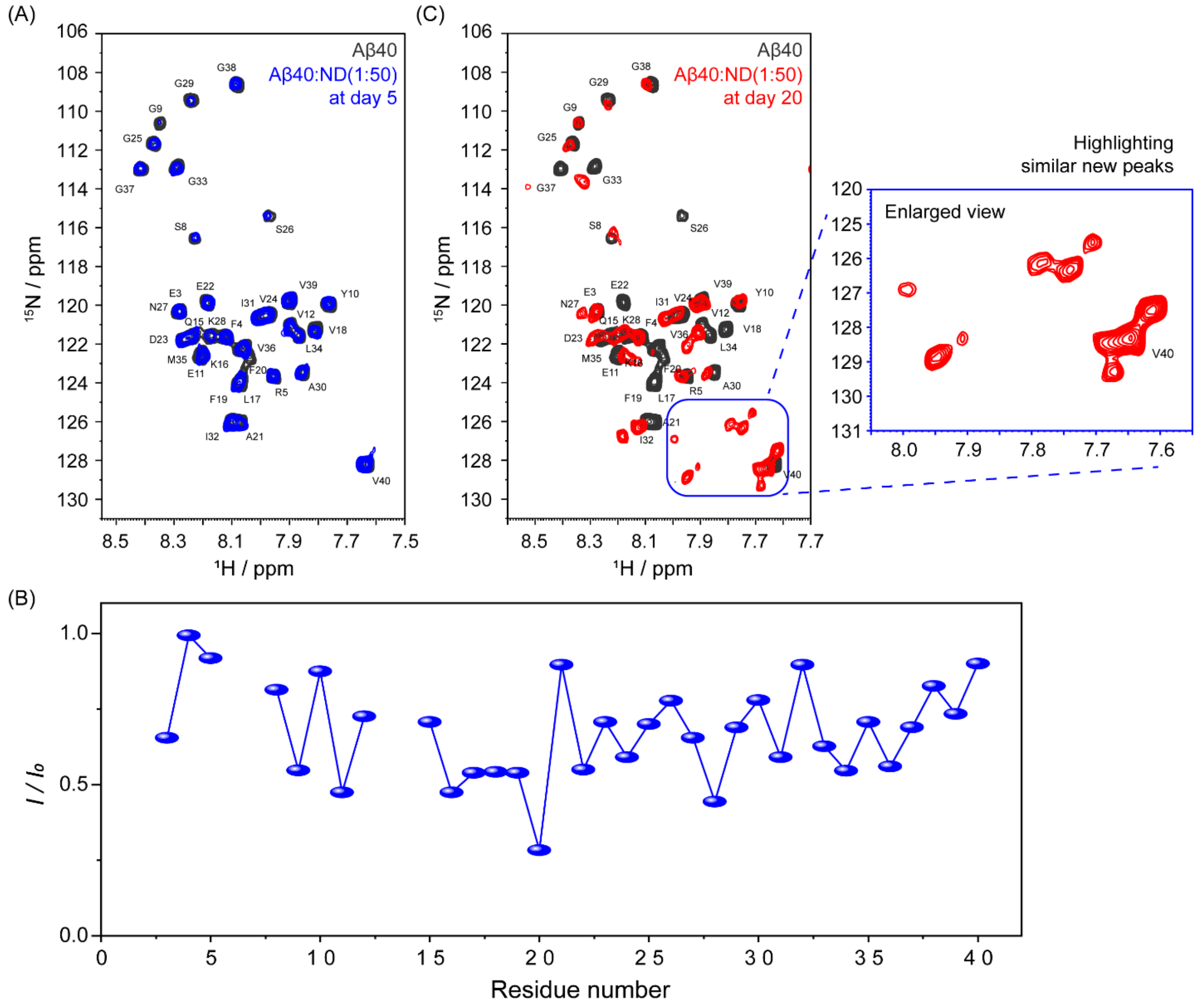


**Figure S24.** (A) Overlay of 2D $^{1}$H-$^{15}$N HSQC spectra of a freshly prepared Aβ40 (80 μM) sample and with nanodiscs (L/P=50) at day 5 indicates only intensity drop and no significance CSPs. (B) Changes in the peak intensities along the peptide backbone measured from 2D $^{1}$H-$^{15}$N HSQC spectra of a freshly prepared Aβ40 (80 μM) sample with nanodiscs (L/P=50) at day 5 indicates involvement of the central hydrophobic region in the interaction. Points represent the signal intensity of the peak maxima (*I(t)*) relative to the signal intensity in the absence of nanodiscs ($I_0$). (C) Overlay of 2D $^{1}$H-$^{15}$N HSQC spectra of a freshly prepared Aβ40 (80 μM) sample and with nanodiscs (L/P=50) at day 20 indicates significant intensity drop and CSPs as well as similar new peaks at the vicinity of V40, thereby suggesting formation of similar oligomeric conformers.

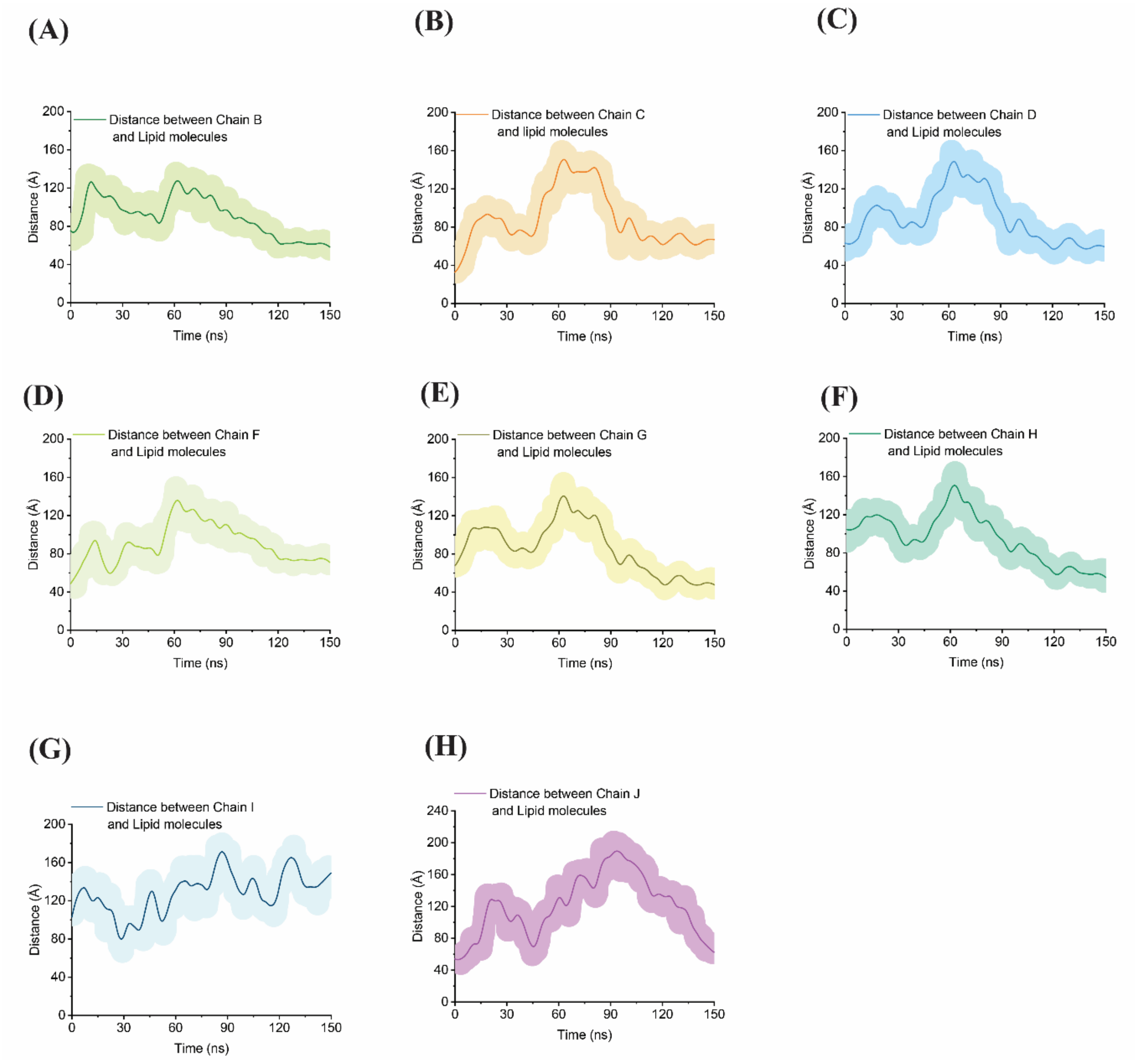


**Figure S25.** Distance between center of mass (COM) of monomers of Aβ40 with COM of lipid molecules during the all-atom (150 ns) MD simulation process.

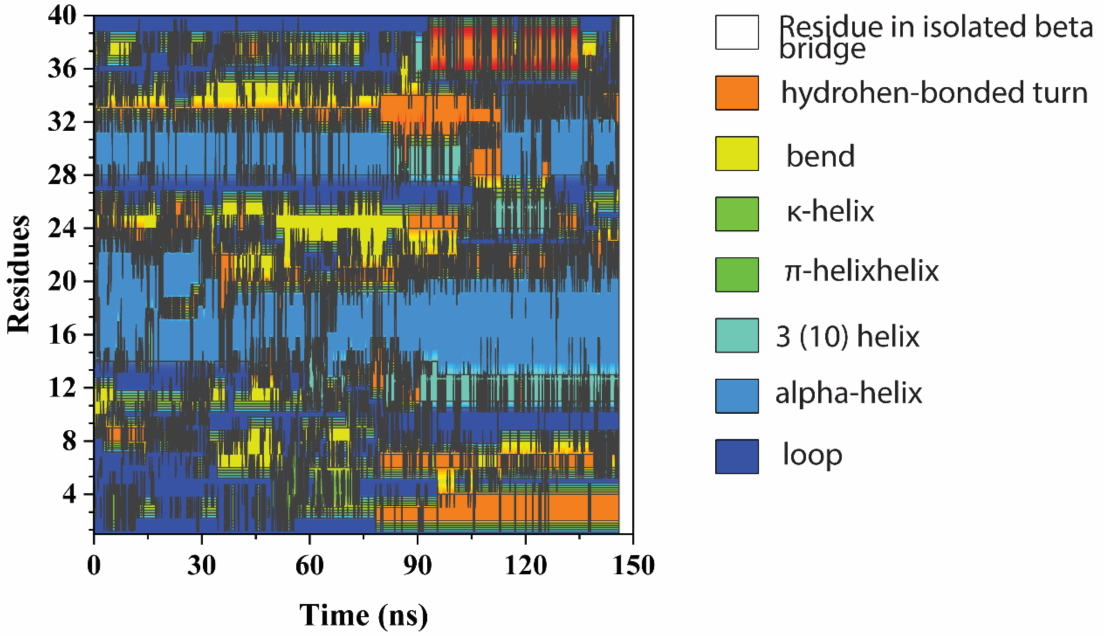


**Figure S26.** DSSP plot showing change in secondary structure of chain E during simulation process.

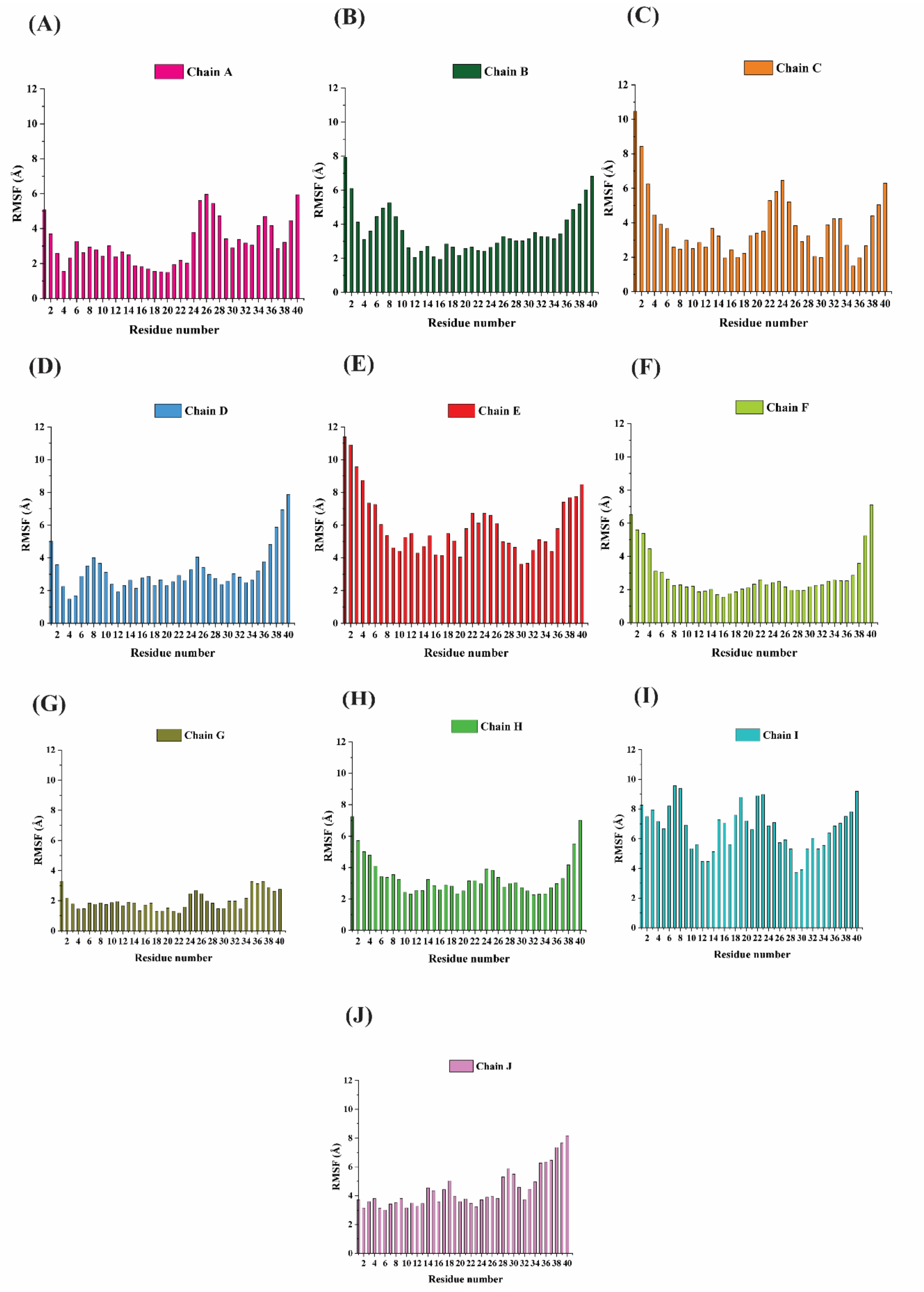


**Figure S27.** Root mean square fluctuation (RMSF) of Aβ40 monomers (Chain A – Chain J) calculated based on backbone of the residues.

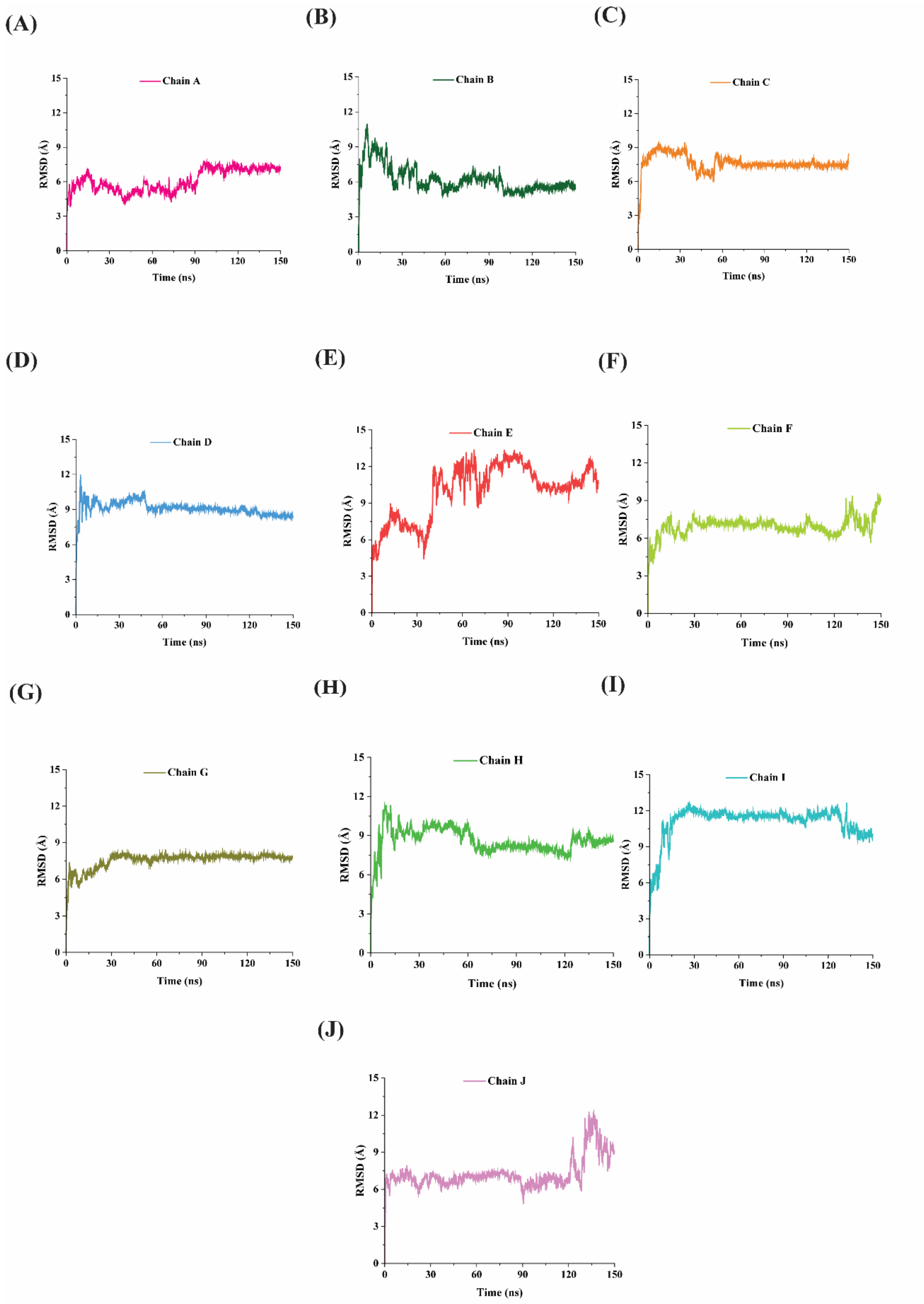


**Figure S28.** Root mean square deviation (RMSD) of Aβ40 monomers (Chain A – Chain J) calculated based on backbone of the residues.

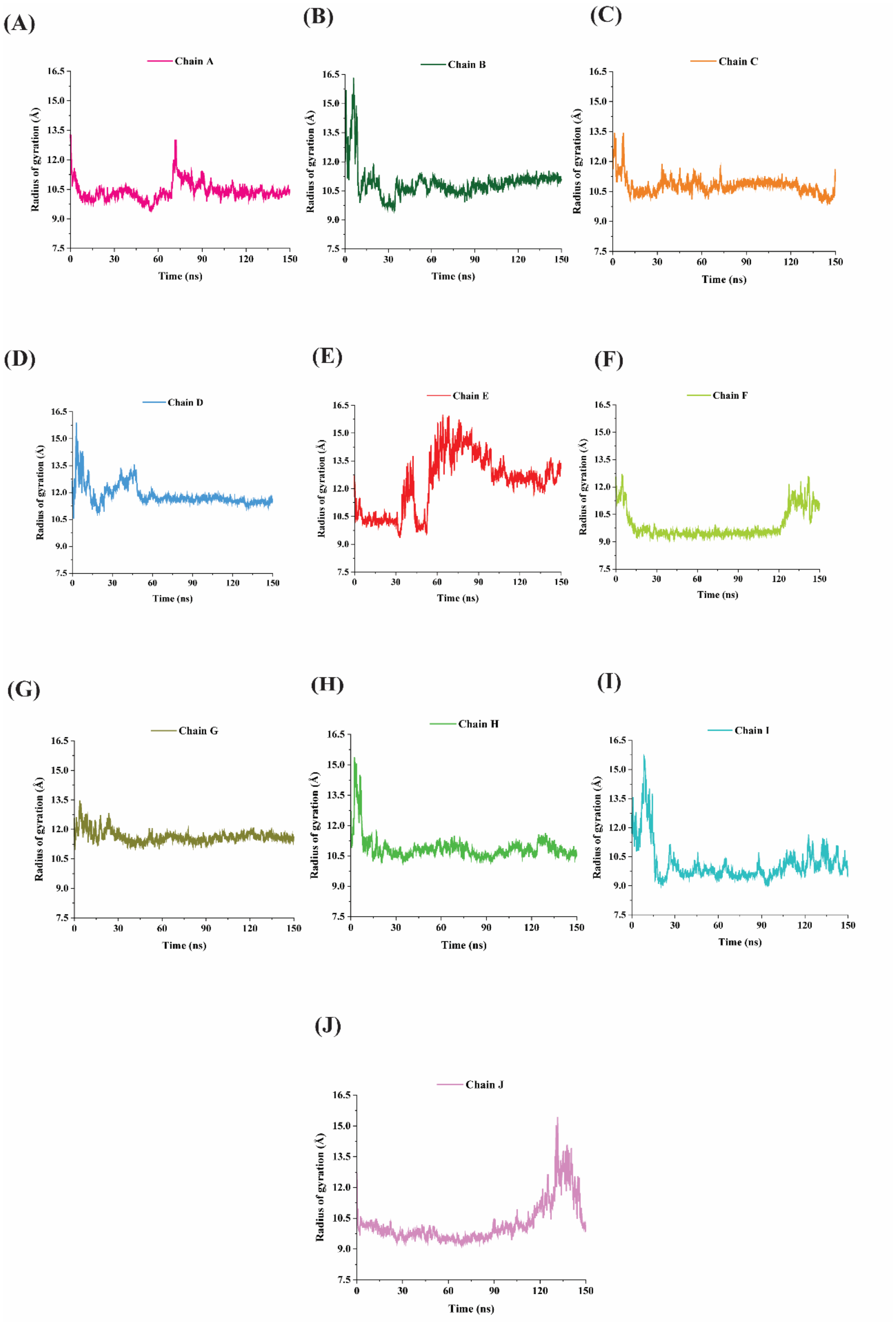


**Figure S29.** Change in radius of gyration of Aβ40 monomers during the simulation process.

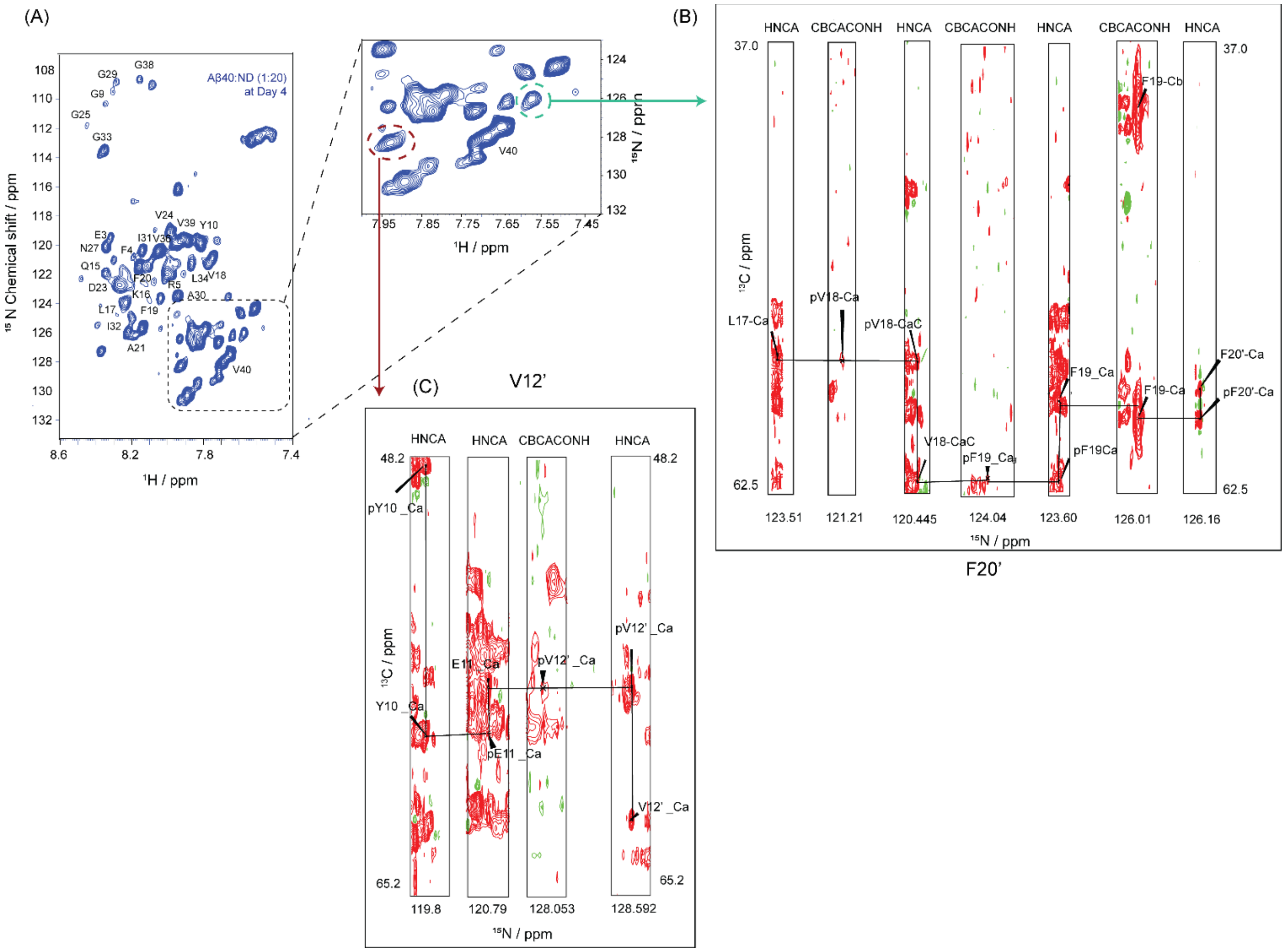


**Figure S30. Characterization of newly developed peaks of AB40 in presence of NDs (L/P=20).** (A) 2D $^{1}H$-$^{15}N$ HSQC spectra Aβ40 (80 μM) sample with NDs (L/P=20) after formation of oligomers at 25 °C. New peaks are shown at inset. Triple resonance (HNCA and CBCACONH) NMR spectra of this sample were utilized to assign the new peaks. (B) The 3D strip of one new peak (cyan color dashed circle) for resonance assignment and this peak is probably new conformational peak of F20, henceforth denoted as F20'. (C) The 3D strip of one new peak (red color dashed circle) for resonance assignment and this peak is probably new conformational peak of V12, henceforth denoted as V12'.

(A)

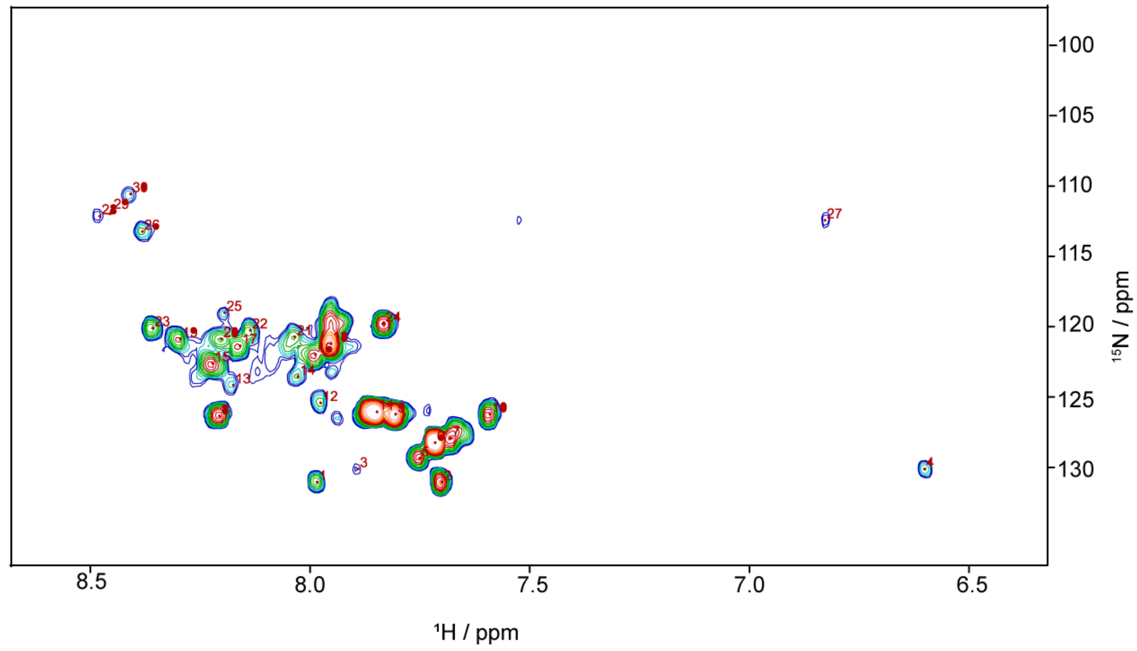


(B)

| Peak name | F1 [ppm] | F2 [ppm] | Kex [1/s ] | error | pE | error | wG | error | wE | error | R1G | error | R2G | error | R2E | error | Io | error | errorScale |
|---|---|---|---|---|---|---|---|---|---|---|---|---|---|---|---|---|---|---|---|
| 1 | 131.0325383 | 7.983256281 | 8.563913311 | 722.3070926 | 0.041860915 | 3.536201735 | 130.7967256 | 0.180874942 | 122.8675802 | 3.481019548 | 0.118920116 | 0.394953216 | 1.409470043 | 11.20547808 | 9.72621106 | 534.5419992 | 1.043421457 | 3.854130139 | 2.144786688 |
| 2 | 131.0325383 | 7.699022705 | 0 | 0 | 0 | 0 | 0 | 0 | 0 | 0 | 0 | 0 | 0 | 0 | 0 | 0 | 0 | 0 | 1 |
| 3 | 130.0946274 | 7.889809626 | 0 | 0 | 0 | 0 | 0 | 0 | 0 | 0 | 0 | 0 | 0 | 0 | 0 | 0 | 0 | 0 | 1 |
| 4 | 130.0946274 | 6.601024508 | 0 | 0 | 0 | 0 | 0 | 0 | 0 | 0 | 0 | 0 | 0 | 0 | 0 | 0 | 0 | 0 | 1 |
| 5 | 129.3130349 | 7.749639644 | 0 | 0 | 0 | 0 | 0 | 0 | 0 | 0 | 0 | 0 | 0 | 0 | 0 | 0 | 0 | 0 | 1 |
| 6 | 128.2188055 | 7.714597148 | 103.9693353 | 324.8030752 | 0.000848047 | 0.002423371 | 128.1819338 | 0.036049397 | 106.9874043 | 1.498531803 | 0.367211696 | 0.04818091 | 0.459730094 | 0.582539782 | 40.840631 | 745.907793 | 1.000996728 | 0.018080105 | 2.144786688 |
| 7 | 127.9061685 | 7.679554652 | 57.61319197 | 1662.546082 | 0.007112528 | 0.079721386 | 128.0991469 | 0.523123951 | 120.9418262 | 6.061449252 | 0.336375067 | 0.258158009 | 0.937823031 | 7.832737618 | 1779.477507 | 12740.30901 | 1.007821265 | 0.103505656 | 2.160368656 |
| 8 | 126.3429837 | 8.205192087 | 0 | 0 | 0 | 0 | 0 | 0 | 0 | 0 | 0 | 0 | 0 | 0 | 0 | 0 | 0 | 0 | 1 |
| 9 | 126.1866652 | 7.804150192 | 0 | 0 | 0 | 0 | 0 | 0 | 0 | 0 | 0 | 0 | 0 | 0 | 0 | 0 | 0 | 0 | 1 |
| 10 | 126.1866652 | 7.590001608 | 0 | 0 | 0 | 0 | 0 | 0 | 0 | 0 | 0 | 0 | 0 | 0 | 0 | 0 | 0 | 0 | 1 |
| 11 | 126.0303467 | 7.846979909 | 1.70228372 | 197.842363 | 0.027778067 | 3.182920458 | 126.0624168 | 0.033000895 | 116.5630487 | 4.325796918 | 0.477215352 | 0.07711607 | 0.872607505 | 1.261640989 | 86.56232631 | 2345.059492 | 1.028541185 | 3.367412991 | 2.144786688 |
| 12 | 125.4050728 | 7.97546906 | 0 | 0 | 0 | 0 | 0 | 0 | 0 | 0 | 0 | 0 | 0 | 0 | 0 | 0 | 0 | 0 | 1 |
| 13 | 124.1545249 | 8.174043202 | 0 | 0 | 0 | 0 | 0 | 0 | 0 | 0 | 0 | 0 | 0 | 0 | 0 | 0 | 0 | 0 | 1 |
| 14 | 123.5292509 | 8.026085998 | 0 | 0 | 0 | 0 | 0 | 0 | 0 | 0 | 0 | 0 | 0 | 0 | 0 | 0 | 0 | 0 | 1 |
| 15 | 122.59134 | 8.22076653 | 23.99243527 | 112839.3039 | 9.39327E-05 | 0.336684528 | 122.3199352 | 2.205221841 | 112.96625 | 1438.474967 | 1.159228979 | 1.271283482 | 6.195946397 | 32.51675643 | 144.8816075 | 8094.216687 | 1.030525013 | 0.541730561 | 2.160368656 |
| 16 | 121.9660661 | 7.987149892 | 239.7086754 | 1677.759607 | 0.000606083 | 0.007831556 | 121.9960858 | 0.161166535 | 131.1987313 | 27.73059579 | 0.68522199 | 0.339301855 | 1.357817581 | 6.475577351 | 1037.633711 | 26591.07423 | 0.995474459 | 0.120445984 | 2.144786688 |
| 17 | 121.3407921 | 8.15846876 | 0 | 0 | 0 | 0 | 0 | 0 | 0 | 0 | 0 | 0 | 0 | 0 | 0 | 0 | 0 | 0 | 1 |
| 18 | 121.1844736 | 7.952107396 | 178.0711864 | 1272.693271 | 0.001368326 | 0.01051271 | 121.184971 | 0.649817252 | 124.4286875 | 9.27621218 | 1.207190894 | 0.087485744 | 0.989948563 | 3.28341326 | 146.2454372 | 2271.061283 | 1.001387624 | 0.031556947 | 2.160368656 |
| 19 | 120.8718367 | 8.294745132 | 0 | 0 | 0 | 0 | 0 | 0 | 0 | 0 | 0 | 0 | 0 | 0 | 0 | 0 | 0 | 0 | 1 |
| 20 | 120.8718367 | 8.201298477 | 0 | 0 | 0 | 0 | 0 | 0 | 0 | 0 | 0 | 0 | 0 | 0 | 0 | 0 | 0 | 0 | 1 |
| 21 | 120.7155182 | 8.03387322 | 0 | 0 | 0 | 0 | 0 | 0 | 0 | 0 | 0 | 0 | 0 | 0 | 0 | 0 | 0 | 0 | 1 |
| 22 | 120.2465627 | 8.135107096 | 0 | 0 | 0 | 0 | 0 | 0 | 0 | 0 | 0 | 0 | 0 | 0 | 0 | 0 | 0 | 0 | 1 |
| 23 | 120.0902442 | 8.357042902 | 0 | 0 | 0 | 0 | 0 | 0 | 0 | 0 | 0 | 0 | 0 | 0 | 0 | 0 | 0 | 0 | 1 |
| 24 | 119.7776072 | 7.831405467 | 127.670503 | 7350.247233 | 0.012590437 | 0.732137566 | 119.942824 | 1.021943587 | 100.6387234 | 279.9533072 | 1.333896234 | 0.392243879 | 0.01150688 | 22.92247503 | 17.70753015 | 18770.08441 | 1.013421301 | 0.75935959 | 2.160368656 |
| 25 | 118.9960148 | 8.193511255 | 0 | 0 | 0 | 0 | 0 | 0 | 0 | 0 | 0 | 0 | 0 | 0 | 0 | 0 | 0 | 0 | 1 |
| 26 | 113.2122308 | 8.380404566 | 0 | 0 | 0 | 0 | 0 | 0 | 0 | 0 | 0 | 0 | 0 | 0 | 0 | 0 | 0 | 0 | 1 |
| 27 | 112.4306384 | 6.826853924 | 0 | 0 | 0 | 0 | 0 | 0 | 0 | 0 | 0 | 0 | 0 | 0 | 0 | 0 | 0 | 0 | 1 |
| 28 | 112.1180014 | 8.477744831 | 0 | 0 | 0 | 0 | 0 | 0 | 0 | 0 | 0 | 0 | 0 | 0 | 0 | 0 | 0 | 0 | 1 |
| 29 | 111.9616829 | 8.454383168 | 0 | 0 | 0 | 0 | 0 | 0 | 0 | 0 | 0 | 0 | 0 | 0 | 0 | 0 | 0 | 0 | 1 |
| 30 | 110.5548166 | 8.40765984 | 0 | 0 | 0 | 0 | 0 | 0 | 0 | 0 | 0 | 0 | 0 | 0 | 0 | 0 | 0 | 0 | 1 |

**Figure S31.** Fitted CEST data obtained from Dynamic Center. (A) Auto assigned peaks in HSQC profile by Dynamic Center on which the CEST NMR was performed. (B) All fitting result obtained directly from Dynamic center, the peak 1 and 16 are highlighted in yellow. Ground state resonance of peak 1 and excited state resonance of peak 16 is marked in green which are closely related. On the other hand, excited state resonance of peak 1 and ground state resonance of peak 16 are marked in blue, this two are also closely related. From here we can say, peak 1 is in chemical exchange with peak 16 or Leu 34 in this case.

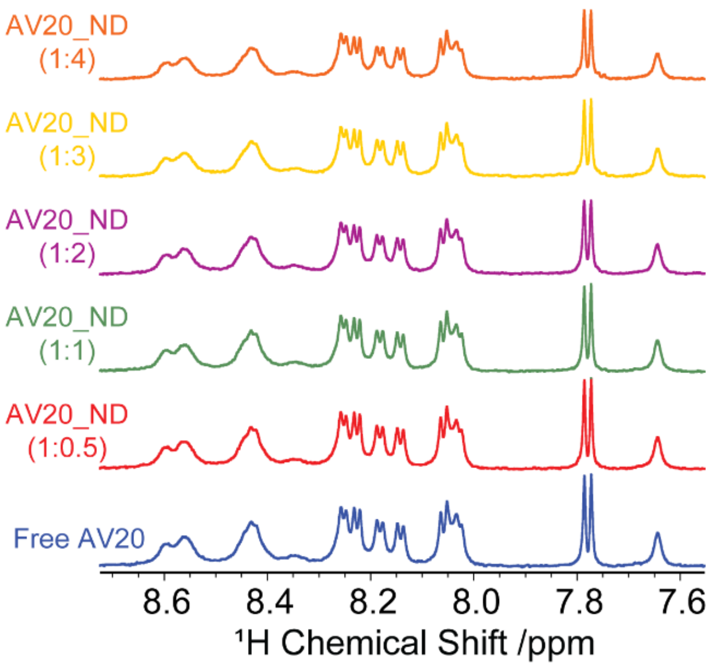


**Figure S32.** Selected 1D $^{1}H$ NMR spectra of AV20 titrated with NDs, showing broadening of the amide proton signal (7.6–8.6 ppm).

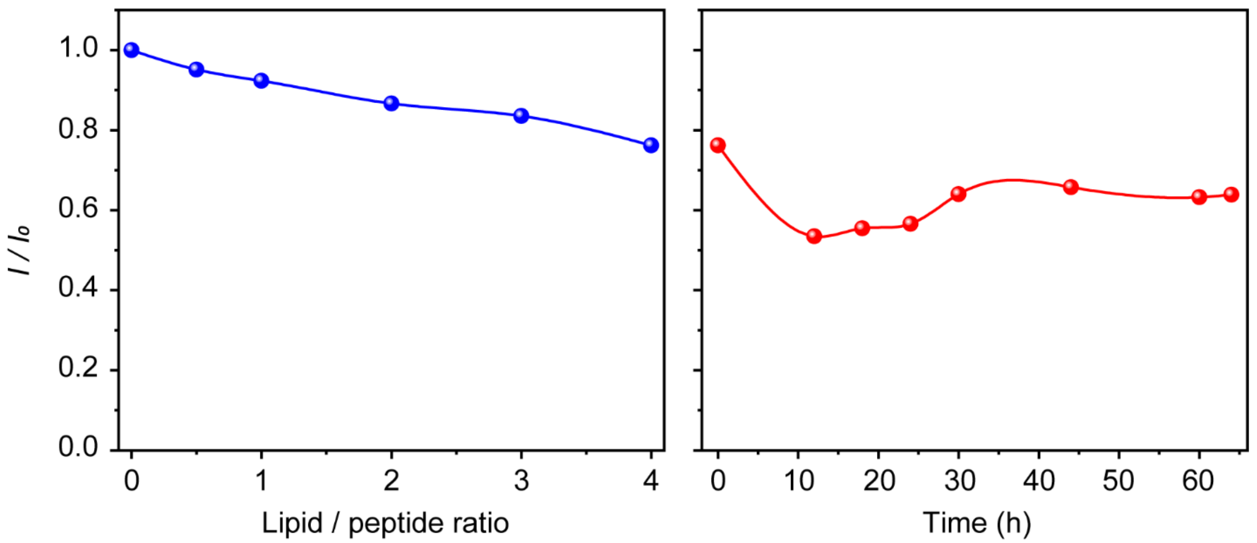


**Figure S33.** Change in amide proton intensity of AV20 with increasing concentrations of NDs (left panel). Time-dependent overall relative intensity plot of AV20 in the presence of NDs (right panel).

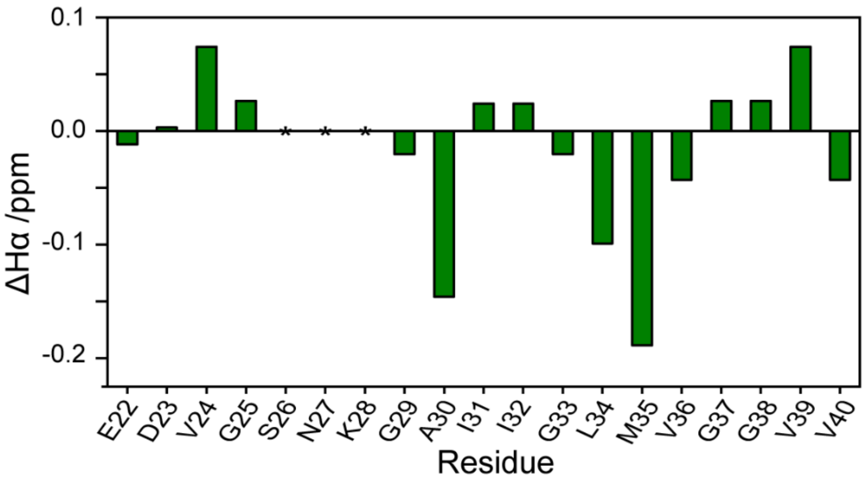


**Figure S34.** The chemical shift index (CSI) was calculated by correcting the Hα from random coil.

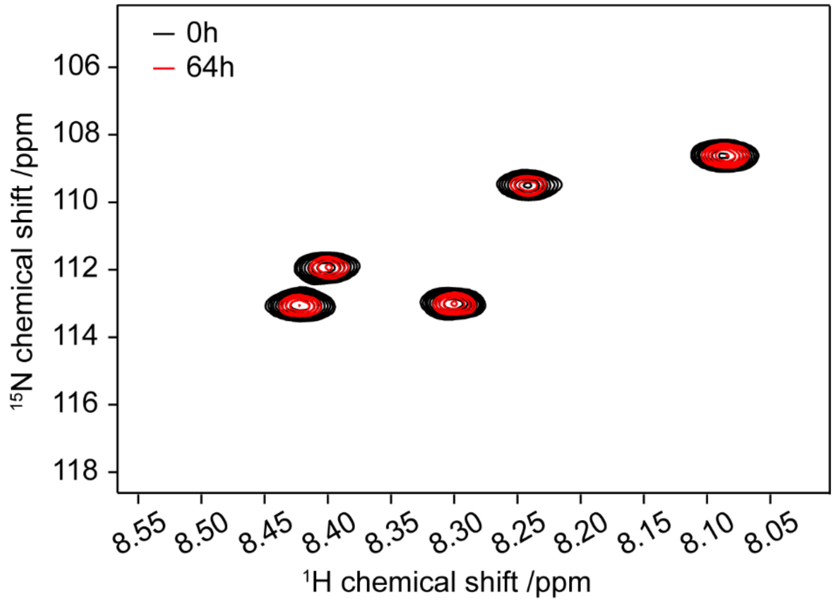


**Figure S35.** Overlay of 2D $^{1}$H-$^{15}$N HSQC spectra of a freshly prepared AV20 (~80 μM) sample at 0 h (black) and at 64 h indicates only intensity drop and no significance CSPs. The sample was incubated at 37 °C and measurements were taken at 10 °C using Bruker 500 MHz NMR spectrometer.

### References

1. Murray, C. J. L., Ikuta, K. S., Sharara, F., Swetschinski, L., Robles Aguilar, G., Gray, A., Han, C., Bisignano, C., Rao, P., Wool, E., Johnson, S. C., Browne, A. J., Chipeta, M. G., Fell, F., Hackett, S., Haines-Woodhouse, G., Kashef Hamadani, B. H., Kumaran, E. A. P., McManigal, B., Achalapong, S., Agarwal, R., Akech, S., Albertson, S., Amuasi, J., Andrews, J., Aravkin, A., Ashley, E., Babin, F.-X., Bailey, F., Baker, S., Basnyat, B., Bekker, A., Bender, R., Berkley, J. A., Bethou, A., Bielicki, J., Boonkasidecha, S., Bukosia, J., Carvalheiro, C., Castañeda-Orjuela, C., Chansamouth, V., Chaurasia, S., Chiurchiù, S., Chowdhury, F., Clotaire Donatien, R., Cook, A. J., Cooper, B., Cressey, T. R., Criollo-Mora, E., Cunningham, M., Darboe, S., Day, N. P. J., De Luca, M., Dokova, K., Dramowski, A., Dunachie, S. J., Duong Bich, T., Eckmanns, T., Eibach, D., Emami, A., Feasey, N., Fisher-Pearson, N., Forrest, K., Garcia, C., Garrett, D., Gastmeier, P., Giref, A. Z., Greer, R. C., Gupta, V., Haller, S., Haselbeck, A., Hay, S. I., Holm, M., Hopkins, S., Hsia, Y., Iregbu, K. C., Jacobs, J., Jarovsky, D., Javanmardi, F., Jenney, A. W. J., Khorana, M., Khusuwan, S., Kissoon, N., Kobeissi, E., Kostyanev, T., Krapp, F., Krumkamp, R., Kumar, A., Kyu, H. H., Lim, C., Lim, K., Limmathurotsakul, D., Loftus, M. J., Lunn, M., Ma, J., Manoharan, A., Marks, F., May, J., Mayxay, M., Mturi, N., Munera-Huertas, T., Musicha, P., Musila, L. A., Mussi-Pinhata, M. M., Naidu, R. N., Nakamura, T., Nanavati, R., Nangia, S., Newton, P., Ngoun, C., Novotney, A., Nwakanma, D., Obiero, C. W., Ochoa, T. J., Olivas-Martinez, A., Olliaro, P., Ooko, E., Ortiz-Brizuela, E., Ounchanum, P., Pak, G. D., Paredes, J. L., Peleg, A. Y., Perrone, C., Phe, T., Phommasone, K., Plakkal, N., Ponce-de-Leon, A., Raad, M., Ramdin, T., Rattanavong, S., Riddell, A., Roberts, T., Robotham, J. V., Roca, A., Rosenthal, V. D., Rudd, K. E., Russell, N., Sader, H. S., Saengchan, W., Schnall, J., Scott, J. A. G., Seekaew, S., Sharland, M., Shivamallappa, M., Sifuentes-Osornio, J., Simpson, A. J., Steenkeste, N., Stewardson, A. J., Stoeva, T., Tasak, N., Thaiprakong, A., Thwaites, G., Tigoi, C., Turner, C., Turner, P., van Doorn, H. R., Velaphi, S., Vongpradith, A., Vongsouvath, M., Vu, H., Walsh, T., Walson, J. L., Waner, S., Wangrangsimakul, T., Wannapinij, P., Wozniak, T., Young Sharma, T. E. M. W., Yu, K. C., Zheng, P., Sartorius, B., Lopez, A. D., Stergachis, A., Moore, C., Dolecek, C., and Naghavi, M. (2022) Global burden of bacterial antimicrobial resistance in 2019: a systematic analysis. *The Lancet* **399**, 629-655
2. Meisl, G., Kirkegaard, J. B., Arosio, P., Michaels, T. C. T., Vendruscolo, M., Dobson, C. M., Linse, S., and Knowles, T. P. J. (2016) Molecular mechanisms of protein aggregation from global fitting of kinetic models. *Nature Protocols* **11**, 252-272
